# Discovery of New CDK8 ligands with a Novel Virtual Drug Screening Tool

**DOI:** 10.1101/321521

**Authors:** Wei Chen, Xiaodong Ren, Chia-en A. Chang

## Abstract

Selective inhibition of CDK8 could be a promising strategy for reducing mitogenic signals in cancer cells with reduced toxic effects on normal cells. As compared with type I ligands, binding of a type II compound often achieves longer residence time. We developed a novel virtual drug screening package which takes advantage of two energy evaluation methods: Superposition and Single-Point Energy Evaluation, and VM2 free energy calculation, and applied it to the discovery of new CDK8 type II ligands. In this research we analyzed binding thermodynamics of 11 published CDK8 type II ligands, and extracted the key binding information to assist virtual drug screening for new ligands. The free energy and MD calculations on the reference CDK8-ligand complexes revealed the important factors in the binding. The urea moiety was found to be the critical structural contributor of the reference ligands. Starting with the urea moiety we implemented virtual drug screening and singled out three compounds for bio-assay testing. The ranking from the experimental result for the three compounds is completely consistent with the predicted rankings by both energy evaluation methods. A potent drug-like compound was discovered to have a K_d_ value of 42.5 nM with CDK8, which is comparable to the most potent reference ligands and provided a good starting point to design and synthesize a series of highly selective and potent CDK8 ligands. Therefore, our novel virtual drug screening package is accurate and efficient enough to be used in drug design projects. We believe this work has significant impact to the field of drug discovery.

## 1. Introduction

Cyclin-dependent kinase 8 and cyclin C (CDK8/CycC) associate with the mediator complex and regulate gene transcription of nearly all RNA polymerase II-dependent genes [1–5]. A number of studies have shown that CDK8/CycC modulates the transcriptional output from distinct transcription factors involved in oncogenic control [6], which include the Wnt/β-catenin pathway, Notch, p53, and TGF-β [7, 8]. Compared with other CDKs, CDK8 is a more gene-specific expression regulator [9, 10] and is differently expressed in cancer [2]. In this view, selective inhibition of CDK8 could be a promising strategy for reducing mitogenic signals in cancer cells with reduced toxic effects on normal cells [11]. The steroidal natural product cortistatin A is the first reported potent and selective ligand of CDK8 [12]. Inhibition of CDK8 with cortistatin A suppresses AML cell growth and has anticancer activity in animal models of AML [13].

The existing ligands fall into two categories based on the major conformations of CDK8 to which they bind. Type I ligands bind to the DMG-in (Aspartate-Methionine-Glycine near the N-terminal region of the activation loop) conformation and occupy the ATP-binding site. The Senexin-type, CCT series, and COT series compounds, which possess 4-aminoquinazoline [14], 3,4,5-trisubstituted pyridine [15] and 6-azabenzothiophene [16] scaffolds, respectively, belong to this category. Senex company identified Senexin B with an IC50 value of 24 nM. The R&D for new type I CDK8 ligands made a significant progress in the past a couple of years and many promising compounds were identified [17, 18]. In 2016 new potent and selective CDK8 ligands with benzylindazole scaffold previously reported as HSP90 ligands were described by Schiemann et al. [19]. One of the most promising molecules showed an IC50 value against CDK8 of 10 nM. Very recently 4,5-dihydrothieno[3’,4’:3,4]benzo[1,2-d] isothiazole derivatives were found to have sub-nanomolar in-vitro potency (IC50: 0.46 nM) against CDK8 and high selectivity [20].

Schneider et al. published the first series of type II CDK8 ligands in 2013 [21]. Type II ligands bind to DMG-out conformation and occupy the ATP-binding site and the allosteric site (deep pocket). The deep pocket is adjacent to the ATP-binding site and is accessible in CDK8 by the rearrangement of the DMG motif from the active (DMG-in) to the inactive state (DMG-out). This pocket is inaccessible in the DMG-in conformation, where the Met174 side-chain is reoriented to make the site available to ATP [22]. Many well-known kinase ligands such as sorafenib and imatinib belong to the type II category. The ligands found by Schneider et al. anchored in the CDK8 deep pocket and extended with diverse functional groups toward the hinge region and the front pocket. These variations can cause the ligands to change from fast to slow binding kinetics, resulting in an improved residence time which is defined as the period for which a protein is occupied by a ligand [23]. As compared with type I ligands, binding of a type II compound to the DMG-out conformation often achieves longer residence time [21, 24]. Residence time is more and more considered to be a key success factor for drug discovery and as important as binding affinity [21].

We were inspired by the discovery of Schneider et al. and furthered the research to discover new type II CDK8 ligands. We investigated the thermodynamics of the binding between CDK8 and existing type II ligands with computational methods. The knowledge gained from this study was then used with our recently developed virtual drug screening package for this ligand discovery effort. This paper is organized as below. We first present the analysis of binding free energies and the energy components of the 11 published CDK8-ligand complexes by Schneider et al. with the help of both free energy calculation and molecular dynamics simulation (detailed in 2.2), from which we extract the key interactions as well as the mechanism in the binding between CDK8 and the ligands. Then we present our virtual drug screening package which is guided by computations and verified by experiments, and the result of its application in the discovery of new type II CDK8 ligands. As a conclusion, we explain the implication of this novel method as a general tool to the drug discovery field.

## 2. Methodologies

### 2.1 Reference CDK8-Ligand Complexes

Among the eleven ligands published by Schneider et al. [21], shown as in Table 1, seven have co-crystal structures with the CDK8 DMG-out conformation. The PDB IDs are 4F6W, 4F7L, 4F7J, 4F70, 4F6U, 4F6S, and 4F7N for ligands **1**, **2**, **3**, **4**, **5**, **7**, and **11**, respectively. The crystal structure 4F6W was employed in all of the energy computations in this research because ligand **1** in this crystal structure has the most extensive interactions with the protein and is one of the most potent ligands in this series. Ligand **1** was also used as the reference ligand in this study for the comparison purpose.

**Table 1.** Structures of the 11 reference ligands

| | R= | PDB ID | $\Delta G(\text{exp})$ | MD (ns) |
| --- | --- | --- | --- | --- |
| 1 |  | 4F6W | -10.324 | 500 |
| 2 |  | 4F7L | -10.979 | 500 |
| 3 |  | 4F7J | -7.184 |  |
| 4 |  | 4F7O | -7.877 |  |
| 5 |  | 4F6U | -8.447 | 500 |
| 6 | H |  | No bind |  |
| 7 |  | 4F6S | -7.533 |  |
| 8 |  |  | -7.965 |  |
| 9 |  |  | -7.482 |  |
| 10 |  |  | -8.078 | 500 |
| 11 |  | 4F7N | -9.739 | 500 |

### 2.2 Methodologies to Investigate the Binding Thermodynamics of Reference Complexes

#### 2.2.1 Free energy Calculation

A rigorous statistical thermodynamics method, called VM2 [25], was used to calculate the binding free energies of CDK8 and its ligands *in silico*. VM2 belongs to a class of methods that focus on the most stable conformations of the molecules, so they are sometimes called predominant states methods. They compute the standard chemical potential of the protein-ligand complex and of the free ligand and protein, and take the difference to obtain the standard free energy of binding,

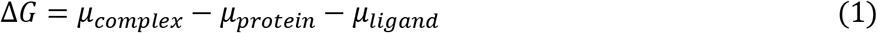

The standard chemical potential of each molecular species (i.e. complex, protein and ligand) is obtained by finding its *M* most stable conformations (*j = 1, M*), integrating the Boltzmann factor within each energy well *j* to obtain a local configurational integral *z _j_*, and combining these local configuration integrals according to the following formula, where *X* = complex, protein, or ligand

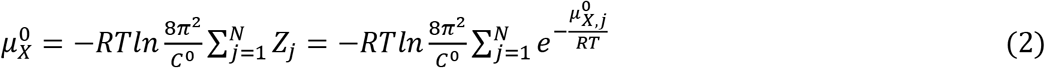

Here *C*^0^ is the standard concentration, which, combined with the factor of 8*π*^2^, accounts for the positional and orientational mobility of the free molecule at standard concentration, and the second form of the summation is given in terms of the chemical potentials of the individual conformations. The probability of energy well j can be approximated on the basis of *z _j_*, and thus the mean potential energy <U> or solvation energy <W> can be obtained. The configurational entropy at standard concentration can be computed as 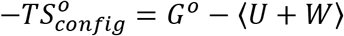. The configurational entropy includes both a conformational part, which reflects the number of energy wells (conformations), and a vibrational part, which reflects the average width of the energy wells.〈 + 〉 implicitly includes the change in solvent entropy via the implicit solvent model. As a consequence, the configurational entropy values reported here should not be directly compared with experimental entropy changes of binding for these systems.

The VM2 calculations are moderately fast, in part because they use implicit solvent models, which are widely accepted as computationally efficient alternatives to more detailed solvent models, and in part because for large systems such as protein-ligand complexes only a subset of atoms (∼500–5000, depending on the nature of the active site) are treated as flexible.

Crystal structure 4F6W was used as a reference structure for VM2 free energy calculations for all of the complexes of CDK8 and the ligands in this study. The starting conformations of the ligands were generated by superimposing them onto ligand 1 in 4F6W. Since 4F6W misses residues 178–195, which form the main portion of the activation loop, a homology model was generated by SWISS_MODEL [26–28] to add the missing residues. Amber99SB and GAFF Force Field (GAFF) [29–31] were applied to CDK8 and the ligands, respectively. The Amber 14 package [32–34] was employed to assign partial charges (-bcc) to the ligands, and the partial charges for the protein were from the standard force field parameters. The live set — the set of binding-site atoms treated as mobile — was defined as all atoms within 7 Å of any atom of the ligands. The real set — the set of protein atoms treated as rigid and thus supporting the live set — comprised all protein residues having any atom within 5 Å of any live-set atom. All other protein atoms were deleted, in order to reduce the size of the non-bonded pair list and thus speed up the calculations. In order to diminish any initial stress in the starting conformations that might artifactually drive the binding site away from its crystallographic starting conformation, the protein models were subjected to an initial relaxation. Van der Waals radii were used in both GB and PB solvation energy calculations. The VM2 method runs cycles of conformational search and configurational integration until convergence criteria is met [25]. The convergence plots for the reference ligands can be found in Figure S1.

#### 2.2.2 Unbiased Molecular Dynamics Simulation

The Amber 14 package with an efficient GPU implementation [32–34] was employed for the MD simulations. Amber 99SB and GAFF were applied to CDK8/CycC and the eleven reference ligands, respectively. Single protonation states were used for all histidine residues according to predictions made by MCCE [35, 36]. Six Cl-ions were placed to maintain a neutral system. Minimization on the hydrogen atoms, side chains and the entire protein complex was performed for 500, 5000, and 5000 steps, respectively, and the system was then solvated with a rectangular TIP3P water box [37] such that the edge of the box is at least 12 Å away from the solutes. The system went through a 1000-step water and 5000-step full-system minimization to correct any inconsistencies. Then we equilibrated the water molecules with the solutes fixed for 20 ns at 298K in an isothermic-isobaric (NPT) ensemble. Next, we relaxed the system by slowly heating it during an equilibrium course of 10 ps at 200, 250 and 298 K. We performed the production run in an NPT ensemble with a 2-fs time step and used the Langevin thermostat [38, 39] with a damping constant of 2 ps^-1^ to maintain a temperature of 298 K. The long-range electrostatic interactions were computed by the particle mesh Ewald method [40] beyond 8 Å distance. The SHAKE algorithm [41] was used to constrain water hydrogen atoms during the MD simulations. We performed 500 ns of MD production runs on each complex and the *apo* protein using CPU parallel processing and local GPU machines. We collected the resulting trajectories every 2 ps and resaved the trajectories for analysis at intervals of 20 ps. The program VMD [42] was used for all 3D molecular graphics in this report.

We performed hydrogen bonding (H bonding) analysis on the MD trajectories. In this study a H-bond (D-H…A, where D and A stand for donor and acceptor respectively) is considered formed if the distance between H and A is smaller than 2.5 Å and the angle of D-H…A is greater than 150º. We used an in-house script to scan the trajectories for direct H bonding between ligands and CDK8. H bonding between ligands and the same residues are merged into one residue-ligand H bonding formation. The occurrence percentage of a hydrogen bond is calculated as the number of the frames where the hydrogen bond is found divided by the total frames (i.e., 25000).

### 2.3 Virtual Drug Screening

We developed a novel structure-based virtual drug screening package to assist the discovery of new drug candidates through efficient and extensive database searches. The workflow of this method is presented in Figure 2. We first computationally study the binding thermodynamics of the existing protein-ligand complexes involving the target protein based on their co-crystal structures. The key moieties on the ligands that make critical contributions to protein binding are identified. Those moieties and their similar structures are used in substructure database searches. In this study the database we used was ChemDiv, which has a collection of over 1,700,000 lead-like, drug-like small molecules in pharmaceutical industry [43]. The compounds obtained after the substructure database searches are then screened with the following criteria:

**Figure 1.**
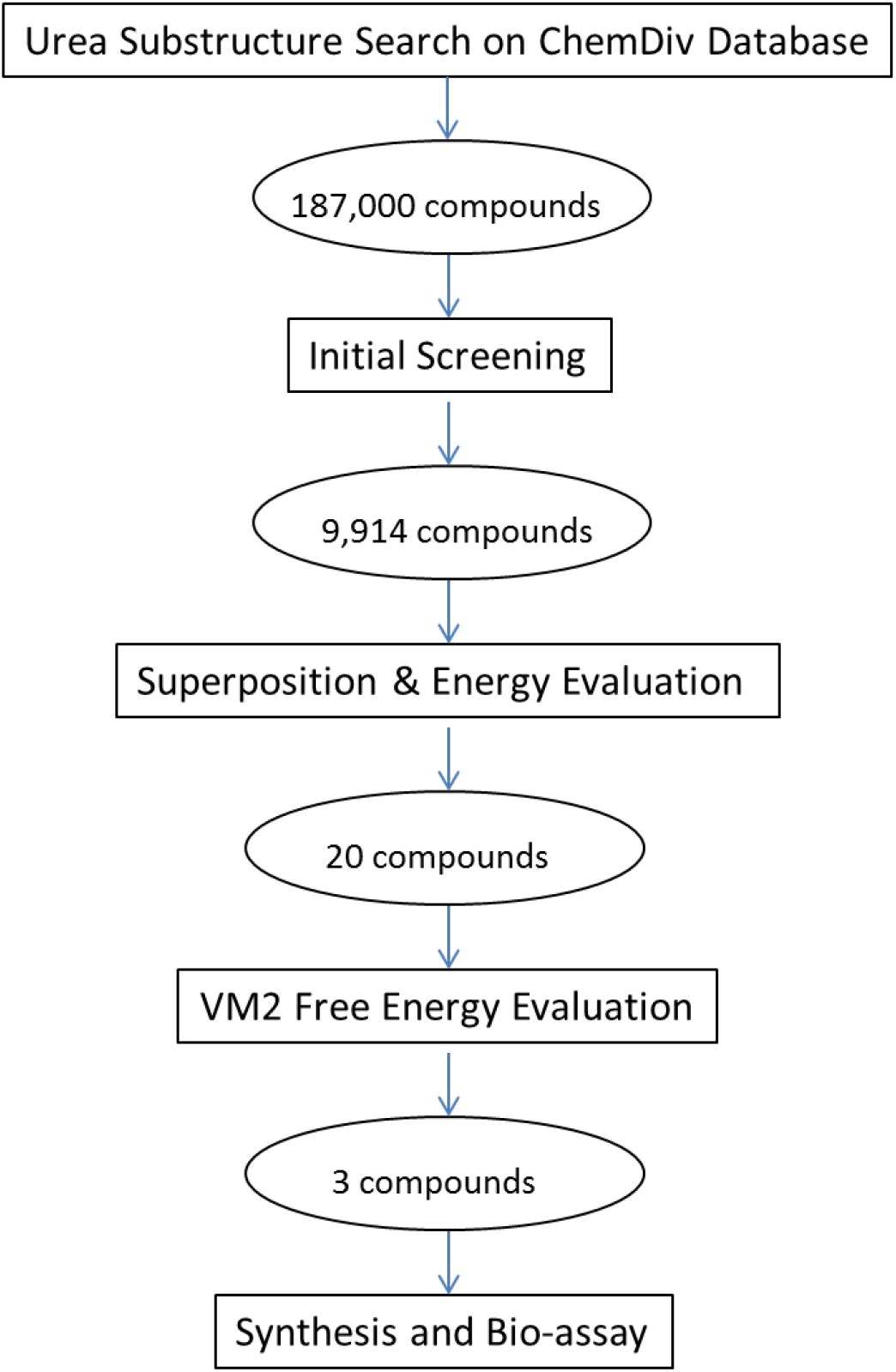
Workflow of the novel virtual drug screening package. The numbers in ovals are the compounds obtained in this study.

**Figure 2.**
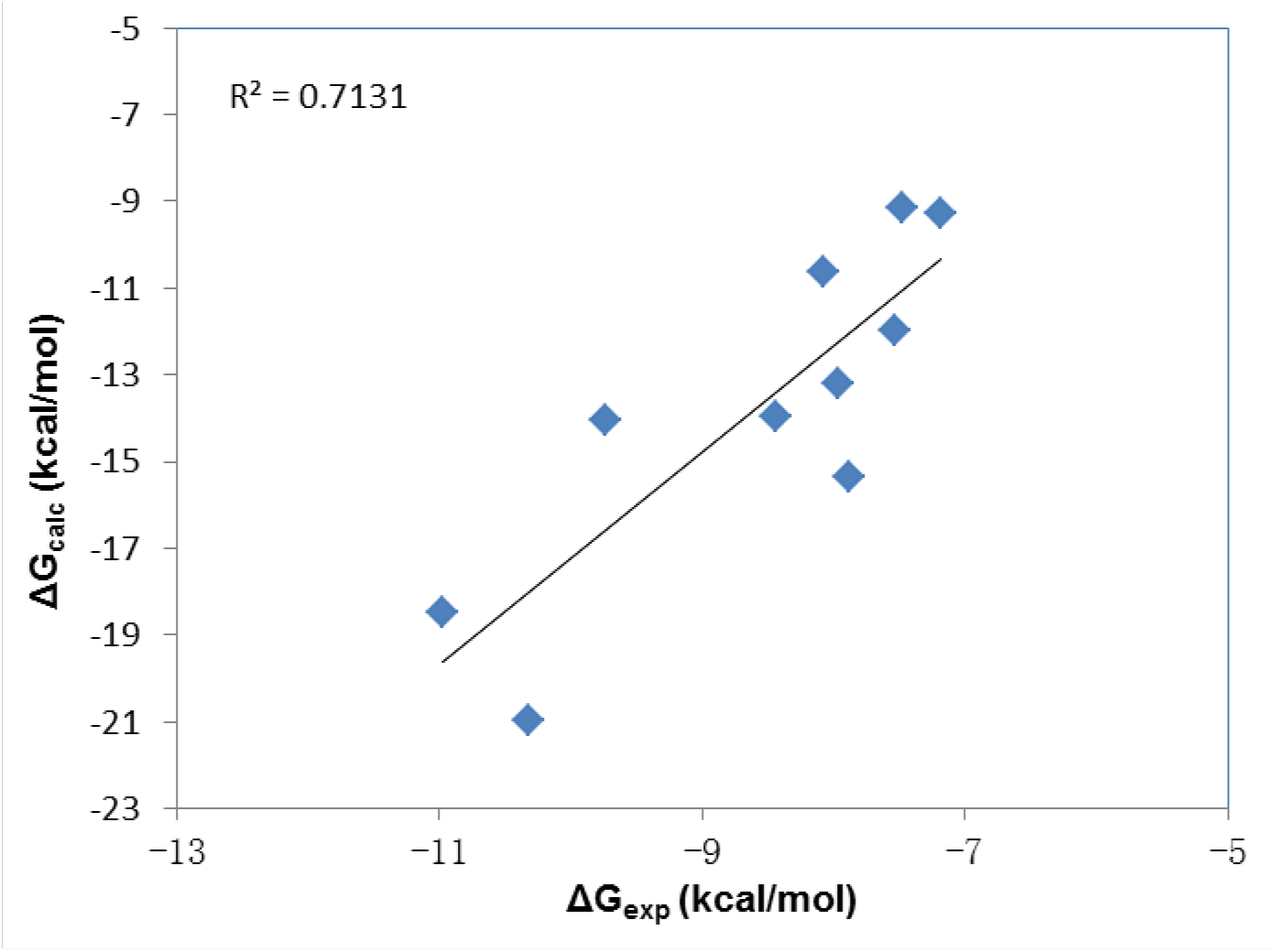
The correlation between the experimental and calculated free Energies for the 11 reference ligands.

1. No more than 5 hydrogen bond donors;
2. No more than 10 hydrogen bond acceptors;
3. A molecular mass less than 600 and greater than 160 daltons;
4. An octanol-water partition coefficient logP less than 5 and greater than –0.4;
5. Specific requirements on the key moieties.

These criteria can be viewed as a modified version of Lipinski’s rule of five [44, 45]. The compounds passing this initial screening are then evaluated and ranked by the Superposition and Single-Point Energy Evaluation method, which is detailed in 2.3.1. The top 20 to 30 compounds from this step are further analyzed with the VM2 free energy calculation method, as described in 2.2.1. The top compounds that have similar or better binding free energies than ligand **1** based on the VM2 evaluation are purchased (or synthesized), and tested with bio-assay to verify the predicted binding affinities.

#### 2.3.1 Superposition and Single-Point Energy Evaluation

Each candidate compound is first superimposed on the conformation of a reference ligand that is at the bound state in a co-crystal structure. In this study ligand **1** in the crystal structure 4F6W was used as the reference ligand. The superposition is implemented by aligning the key moieties of the candidate compound and ligand **1**. In the case that there are more than one key moieties, superposition is implemented with one moiety at a time; if a moiety is symmetric, the candidate compound is rotated to generate all possible superimposed conformations.

After a candidate compound is placed into the binding site of a target protein with possibly multiple conformations, these conformations of the candidate compound together with the target protein are then energy minimized using the Amber 14 package until convergence. A live set is defined similarly as in the VM2 method. Only the compound atoms and the protein atoms in the live set are relaxed in energy minimization. The energy of the complex *E_complex_* is calculated with the equation

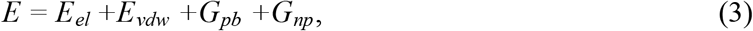

where *E_el_* and *E_vdw_* are electrostatic and van der Waals energy with parameters of Amber99SB for the protein and GAFF for the ligands; *G_pb_* is the solvation energy computed by solving the Poisson Boltzmann (PB) equation [46, 47]; *G_np_* is the nonpolar energy estimated from solvent accessible surface area.

The protein and the candidate compound are then separated from the complex, but their conformations are kept fixed. Their energies, *E_protein_* and *E_ligand_*, are calculated by equation (3). The binding energy for the compound is then calculated as

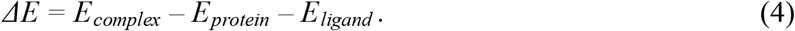

Theoretically this Superposition and Single-Point Energy Evaluation method is faster and more accurate than the conventional docking and scoring because of the following reasons. Firstly, the binding conformations of the candidate molecules are obtained by direct superposition to the available crystal structure of a reference molecule. It thus saves time of generating millions of random conformations as the general practice in docking and still has more accurate binding mode with the protein. Secondly, the binding energy includes the solvation energy, and is calculated with force-field based energy equations and the PBSA model, thus making more physical sense than scoring functions. Therefore, it is quite accurate and efficient for screening a large number of candidates. However, this method doesn’t include entropy contribution and only accounts for the energy contribution from a single conformation. The top compounds obtained from this method are further analyzed and ranked using the more accurate VM2 free energy calculation method.

### 2.4 Compounds and Bio-assay

In this study the compounds selected for bio-assay testing were purchased from ChemDiv Inc. Bio-assay testing was conducted by Proteros biostructures GmbH [48]. The Proteros Reporter Displacement Assay was used to determine K_d_. Briefly speaking, the Proteros reporter displacement assay is based on reporter probes that are designed to bind to the site of interest of the target protein. The proximity between reporter and protein results in the emission of an optical signal. Compounds that bind to the same site as the reporter probe displace the probe, causing signal diminution. Reporter displacement is measured over time after addition of compounds at various concentrations. In order to ensure that the rate of probe displacement reflects compound binding and not probe dissociation, probes are designed to have fast dissociation rates. Thus, compound binding and not probe dissociation is the rate limiting step of probe displacement. For K_d_ determination, percent probe displacement values are calculated for the last time point, at which the system has reached equilibrium. For each compound concentration, percent probe displacement values are calculated and plotted against the compound concentration.

## 3. Results and Discussion

### 3.1 Analysis on binding thermodynamics of the reference complexs

#### 3.1.1 Overall analysis on binding free energies and conformations

Computed and measured binding free energies for the reference CDK8 type-II ligands are compared and plotted in Figure 2, and the corresponding free energy components are presented in Table 2. A significant correlation is observed, as indicated by the correlation coefficient of 0.71, which is comparable to those obtained for other protein-ligand systems via the VM2 method [25]. According to the free energy components in Table 2, it appears that the major driving force for the bindings of all the reference ligands is the van der Waals interactions. Ligand **6** has the least van der Waal interaction with the protein. As a result, its calculated ΔG is –6.45 kcal, which is equivalent to 20 μM, consistent with the experimental result of no binding. The overall electrostatic interactions (Coulomb + desolvation) provide a negative contribution to the binding. The entropy penalties correctly reflect the flexibility of the ligands. For example, the variable substituents are linear and become longer on ligands **6, 7**, **8**, **3**, **9**, **10**, **11**, and their entropy penalties become larger in the same order. The same phenomenon is also observed on ligands **4** and **5**. It is noteworthy that ligands **1** and **2** have the strongest binding affinities with CDK8 among the 11 ligands but pay above-average entropy penalties.

**Table 2.** Detailed energy breakdowns of computed CDK8-ligands binding free energies ΔG_calc_, along with experimental binding free energies, ΔG_exp_. Change in mean energy associated with force-field bond-stretch, angle-bend and dihedral terms (ΔE_Valence_); change in mean force-field Coulombic energy (ΔE_Coulomb_); change in mean Poisson-Boltzmann solvation energy (ΔE_PB_); change in mean nonpolar surface energy (ΔE_NP_); change in mean force-field Lennard-Jones energy (ΔE_VDW_); change in mean total energy ΔH, the sum of the prior five terms; and change in configurational entropy contribution to the free energy (-TΔS). Unit is kcal/mol. RMSD: root-mean-square deviation of most stable computed conformation of the complex for ligand alone/all mobile atoms (Angstroms).

| | $\Delta E_{\text{Valence}}$ | $\Delta E_{\text{Coulomb}}$ | $\Delta E_{\text{PB}}$ | $\Delta E_{\text{NP}}$ | $\Delta E_{\text{VDW}}$ | $\Delta H$ | $-T\Delta S$ | $\Delta G_{\text{calc}}$ | $\Delta G_{\text{exp}}$ | RMSD |
| --- | --- | --- | --- | --- | --- | --- | --- | --- | --- | --- |
| 1 | 1.031 | -19.845 | 60.764 | -8.548 | -93.837 | -60.435 | 39.487 | -20.948 | -10.324 | 1.457/1.753 |
| 2 | 1.498 | -15.235 | 39.754 | -6.577 | -68.517 | -49.077 | 30.610 | -18.467 | -10.979 | 1.810/0.728 |
| 3 | 2.338 | -20.051 | 36.290 | -4.754 | -52.065 | -38.242 | 28.983 | -9.259 | -7.184 | 1.727/0.600 |
| 4 | 2.217 | -16.014 | 41.236 | -5.760 | -63.637 | -41.958 | 26.605 | -15.353 | -7.877 | 1.637/1.318 |
| 5 | 2.065 | -14.742 | 38.291 | -5.760 | -62.614 | -42.760 | 28.792 | -13.968 | -8.447 | 1.851/2.492 |
| 6 | 2.512 | -6.755 | 25.610 | -3.652 | -45.295 | -27.580 | 21.130 | -6.450 | No bind |  |
| 7 | 1.448 | -14.167 | 31.055 | -4.049 | -49.417 | -35.130 | 23.157 | -11.973 | -7.533 | 1.841/0.414 |
| 8 | 2.911 | -16.977 | 33.393 | -4.265 | -53.090 | -38.028 | 24.416 | -13.612 | -7.965 |  |
| 9 | 2.409 | -21.251 | 37.597 | -4.978 | -54.164 | -40.387 | 31.244 | -9.143 | -7.482 |  |
| 10 | 4.992 | -19.615 | 39.659 | -5.191 | -62.883 | -43.038 | 32.392 | -10.646 | -8.078 |  |
| 11 | 1.566 | -29.122 | 43.449 | -5.654 | -58.240 | -48.001 | 33.962 | -14.039 | -9.739 | 1.675/0.318 |

Figure 3 provides a sense for the conformational variation among the 11 bound ligands via an overlay of their most stable predicted conformations. The common scaffold which occupies the region called “deep pocket” takes a uniform pose, while, not surprisingly, the variable substituents which occupy the region called “front pocket” show a wider range of positions. RMSDs for all the mobile atoms (i.e., live protein atoms plus ligand atoms) are quite similar and all less than 2.0 Å, indicating a good matching between the predicted poses and the crystal structures. Ligand **1** has the lowest RMSD for all the mobile atoms presumably because we used the protein in 4F6W, which is the co-crystal structure of the protein and ligand **1**, in the VM2 calculations for all the ligands while the RMSD was calculated between a ligand together with the protein and its available crystal structure. The ligands with RMSDs for ligand alone less than 1.0 Å have nearly identical poses as their crystal structures. For the ligands with larger RMSDs for ligand alone, a portion of their structures deviate from the crystal structures. Ligand **5** has the largest RMSD for ligand alone because in the predicted pose its morpholine moiety moves away from A100 and thus loses the H bonding with this residue. Similarly the morpholine ring in ligand **4** deviates from its crystal counterpart as well. For ligand **1**, it is the piperazine ring that has the large deviation.

**Figure 3.**
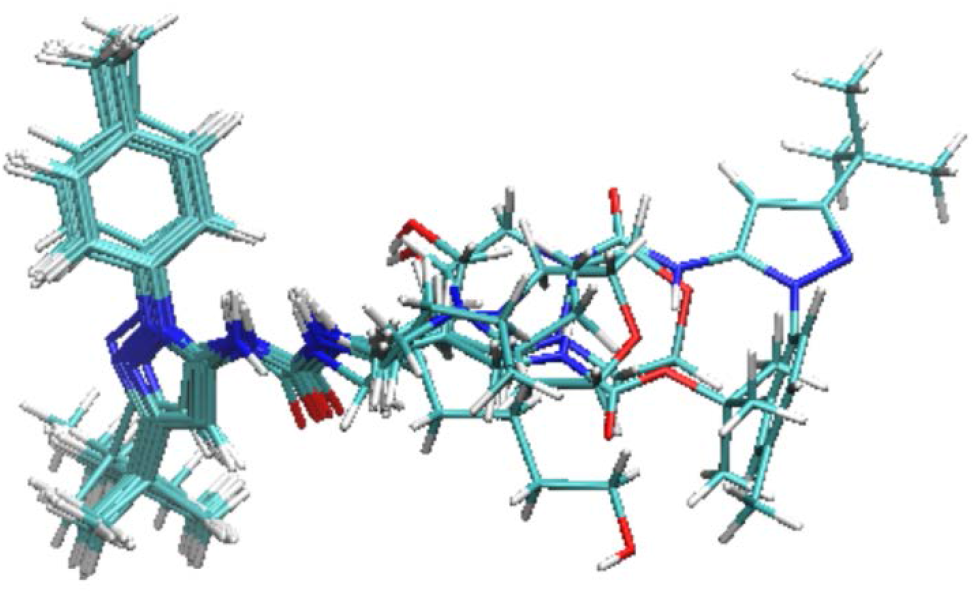
The overlay of the most stable conformations of the 11 reference ligands in the binding pocket of CDK8. The protein is not shown for clarity.

We plotted the relationship between ligand sizes (i.e., the number of non-hydrogen atoms) and the experimental free energies, because our free energy calculation shows that the binding is mainly driven by van der Waals interaction and ligand sizes have strong correlation with van der Waals interactions. Many scoring functions use ligand sizes to assess ligands [49]. The plot can be found in the supplement materials (Figure S1). The correlation coefficient is 0.48. Interestingly the data points on the plot have very similar distribution as those in Figure 1. This result suggests that even for the congeneric ligand series the driving force of which is van der Waals interaction other factors such as electrostatic interaction and entropy are still important to rank ligands accurately. It is thus of interest to examine the relationship between ΔH, which includes not only van der Waals but also all other energy terms except entropy, and the experimental free energies. The resulting correlation coefficient is 0.61 and again the data points on the plot have very similar distribution as those on the two above-mentioned plots (refer to Figure S2). The new correlation coefficient is lower than the one for the full computed free energies (0.71) by 15%, echoing the results in the previous studies [25]. This result highlights the importance of the configurational entropy to the correlation between calculation and experiment.

We calculated the MMPBSA energies for ligands **1**, **2**, **5**, **10** and **11** with the Amber package, and plotted the relationship between the MMPBSA energies and the experimental free energies as well. The correlation coefficient is poorly 0.4, much worse than the correlation coefficient of 0.72 when only the same five ligands are considered in the VM2 plot. The MMPBSA method is also an end-point method that uses a force-field and an implicit solvent model to estimate binding free energies. A key difference from the VM2 method is that MMPBSA calculations typically either neglect configurational entropy or approximate it as average vibrational entropy over essentially randomly selected, energy-minimized molecular dynamics snapshots. Furthermore, the MMPBSA method samples conformations by molecular dynamics, which is far less thorough than the aggressive conformational searches in the VM2 method. It is thus not surprising that the MMPBSA method had a much worse performance than the VM2 method.

#### 3.1.2 Close-up analysis on selected reference complexes

Figure 4 presents the major interactions in the predicted conformation of CDK8 and ligand **1**. The residues E66 and D173 form strong H bonding with the urea linker on **1**, and K52 has a salt bridge with E66. R356 has a cation-π interaction with the benzene ring on **1**. All of these interactions can be found in the crystal structure 4F6W as well. Ligand **1** has strong van der Waals interaction with the protein. The free energy decomposition result shows that residues Y32, E66, D173, M174, R356 and F97 contribute the most to van der Waals energy. Ligand **1** pays the largest entropy penalty among the reference ligands, presumably due to its large yet relatively flexible structure. In addition, ligand **1** pays much larger desolvation penalty than other ligands but its Columbic energy is only on the average level, resulting in the poorest overall electrostatic energy. This indicates that many polar groups on ligand **1** don’t contribute to binding.

**Figure 4.**
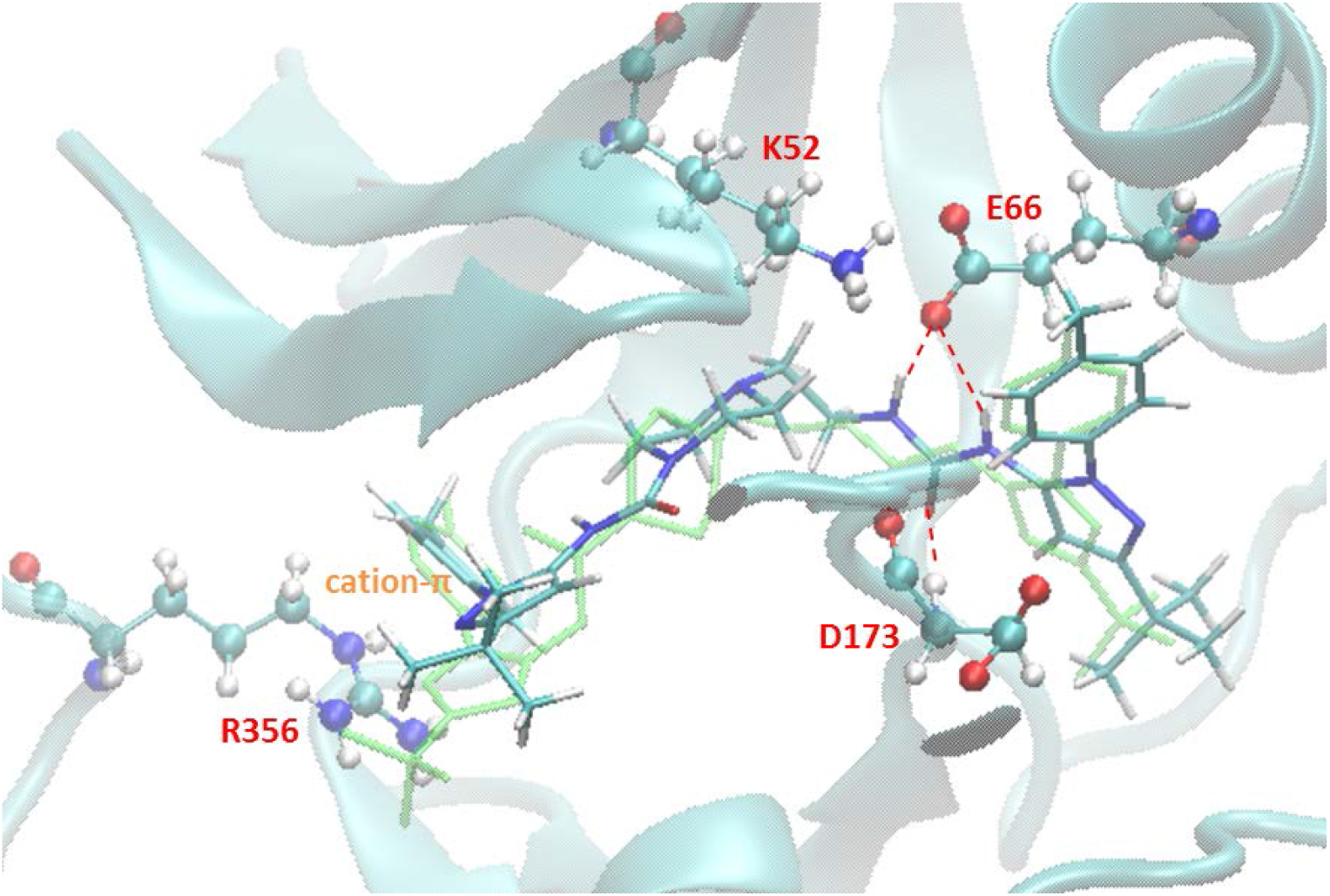
Major interactions found in the predicted lowest energy conformation of CDK8 and ligand **1**.

Ligand **2** has similar binding mode as **1**, except the cation-*π* interaction with R356. Instead, its t-butyl group interacts with R356 through van der Waals interaction. Schneider et al. thought this interaction plays an important role in the binding of ligand **2** [21]. However, according to the free energy decomposition result residues E66, D173, L70 and F97 contribute the largest van der Waals energy to the binding. The contribution from R356 is relatively small. Our calculation shows that the strong affinity of **2** comes from the well-balanced energy terms. It has the second largest molecular size and van der Waals energy; it pays relatively small entropy penalty; its desolvation penalty is much smaller than that of **1** and on the same level as those of other strong binders such as ligands **5**, **10** and **11**.

As discussed in 3.1.1, ligand **5** has the largest RMSD for ligand alone among all the reference ligands. In the most stable predicted conformation, the morpholine ring loses the H bonding interaction with A100 which is assumed to be important to the binding of ligand **5** to CDK8 [21]. As a result, its calculated Coulombic energy is relatively low, which causes the underestimation of its computed binding free energy. The MD simulation didn’t catch this H bonding interaction either. It is possible that the parameters of GAFF force field for morpholine need to be tweaked to pick up this H bonding. The major van der Waals contributors of **5** are E66, D173, L70, A172 and M174. But the total van der Waals energy is much weaker than those of **1** and **2**.

The VM2 method accurately predicted the binding pose of ligand **11**. Its RMSD for ligand alone is the smallest among the reference ligands. Its hydroxyl group forms strong H bonding interactions with D98 and A100. Probably for this reason it achieves the strongest Coulombic energy and the most favorable overall electrostatic energy. Its van der Waals energy is much smaller than those of other strong binders though, and it pays large entropy penalty due to its long and flexible carbon chain. Therefore Coulombic energy plays an important role in the binding of ligand **11** to CDK8. It is the perfect length of the carbon chain that helps the strong binding to occur when compared with ligands **3**, **9** and **10**.

### 3.2 Key information learned from the computational binding thermodynamics study

The VM2 method predicted binding free energies with high correction with experimental data for the reference ligands. It also revealed information that is not available from experiments. The driving force for the binding of the CDK8 type-II ligands is van der Waals interaction, and the overall electrostatic interaction has a negative contribution to the binding affinity. The analysis on the strong binders suggests that there is room for all of them to improve their binding affinities. Specifically, ligand **1** pays extremely large desolvation and entropy penalties. This can be improved by making the structure less flexible and position the polar groups at places that promote H bonding. Ligands **2** and **5** have poor overall electrostatic energy. Polar groups can be introduced to form H bonding with the protein to improve the electrostatic energy. Ligand **11** pays large entropy penalty. Its linear carbon chain can be modified to a more rigid structure to reduce the entropy penalty.

On the predicted lowest energy conformations of all the reference ligands, the urea moiety forms H bonding with residues E66 and D173, and accounts for the majority of H bonding between the protein and the ligands. The 500 ns MD simulations show that these hydrogen bonds are highly stable. Their occurrence percentages are roughly 90∼96% and 76∼93% respectively for all the reference ligands. Therefore, we used the urea moiety to initiate our virtual drug screening research for new type II CDK8 ligands.

### 3.3 Virtual Drug Screening

As the first step the urea moiety was used in the substructure search on ChemDiv database. This resulted in 187,000 compounds (Figure 1). These compounds were screened with the criteria detailed in section 2.1 with a moiety specific requirement: each nitrogen atom of the urea moiety must have one attached hydrogen atom. These hydrogen atoms are needed in H bonding with E66. After this initial screening step, the group of candidates is reduced to 9,914 compounds.

The reduced group was then processed with the Superposition and Single-Point Energy Evaluation method. It took 5 to 10 minutes to process one compound on one core with a Xeon E5–2640 v2 @ 2.00GHz CPU. With 20 cores we finished this energy evaluation step in two days. The binding energies of top 100 compounds are listed in Table S1, and the molecular structures of top 20 compounds are listed in Table S2. Top 20 compounds have highly diverse molecular structures. We also applied this evaluation method on ligand 1, and found it has a lower binding energy than all of the candidates. After superposition and energy minimization, all of them have the similar binding mode as ligand **1**. Figure 5 presents the conformations of ligand **1** and the compound ranked #1 (called compound 1 hereafter) by the Superposition and Single-Point Energy Evaluation method. The predicted conformation of ligand **1** can be aligned with the crystal structure 4F6W very well. For compound 1, besides the key interactions with E66 and D173 it also forms a hydrogen bond with A100, a residue in the hinge region which is important to ligand binding and residence time [21]. The predicted conformations of compounds 2, 3, 4 and 5 with CDK8 are demonstrated in Figure S3. Compounds 2 and 4 have the same binding mode as compound 1, interacting with E66, D173 and A100 through hydrogen bonds. Compound 3 doesn’t have the interaction with A100, but its 1,3,4-thiadiazole forms H bonding with K52. Compound 5 doesn’t have contact with A100 either. Instead its indole ring forms another H bonding with D173.

**Figure 5.**
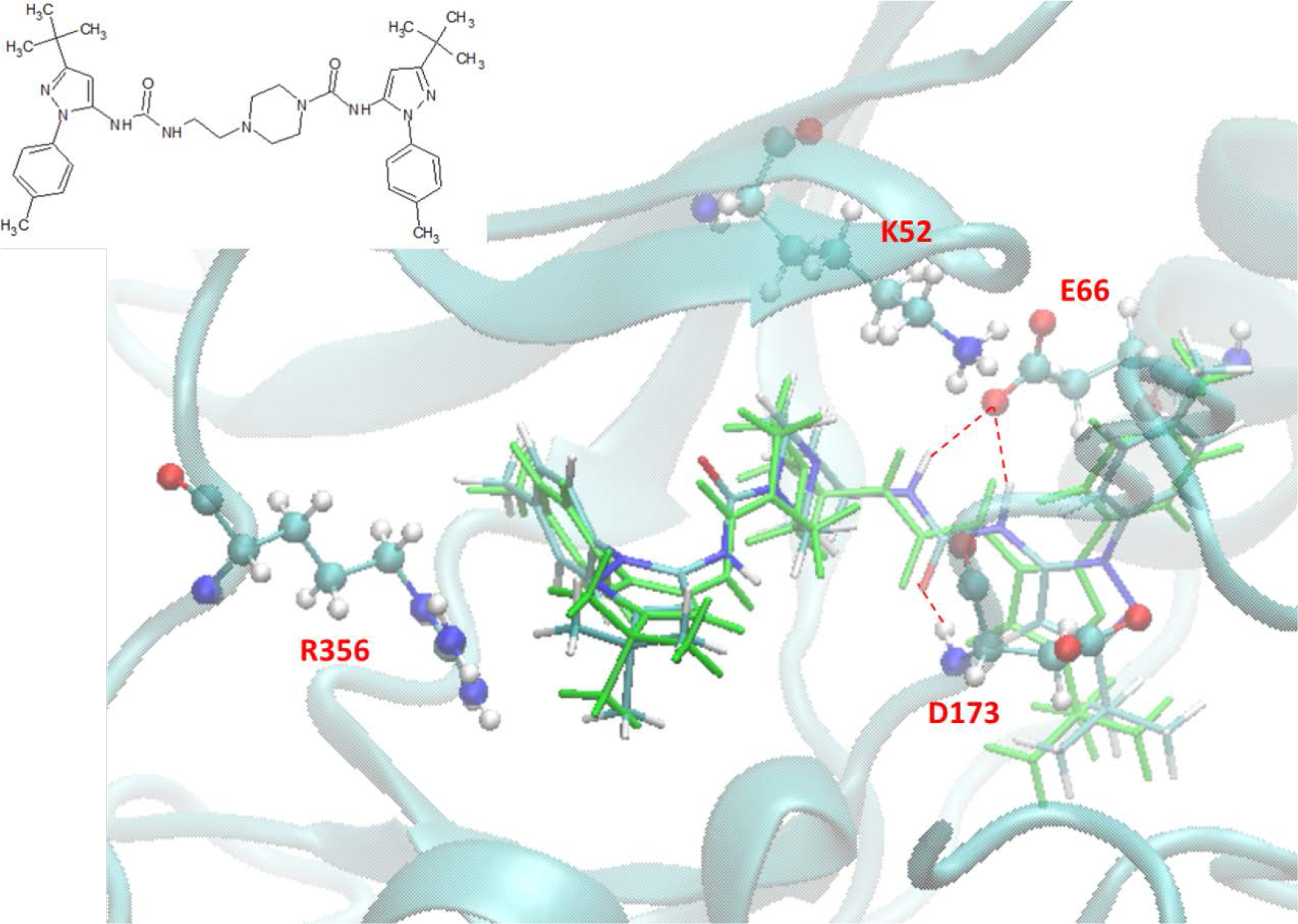
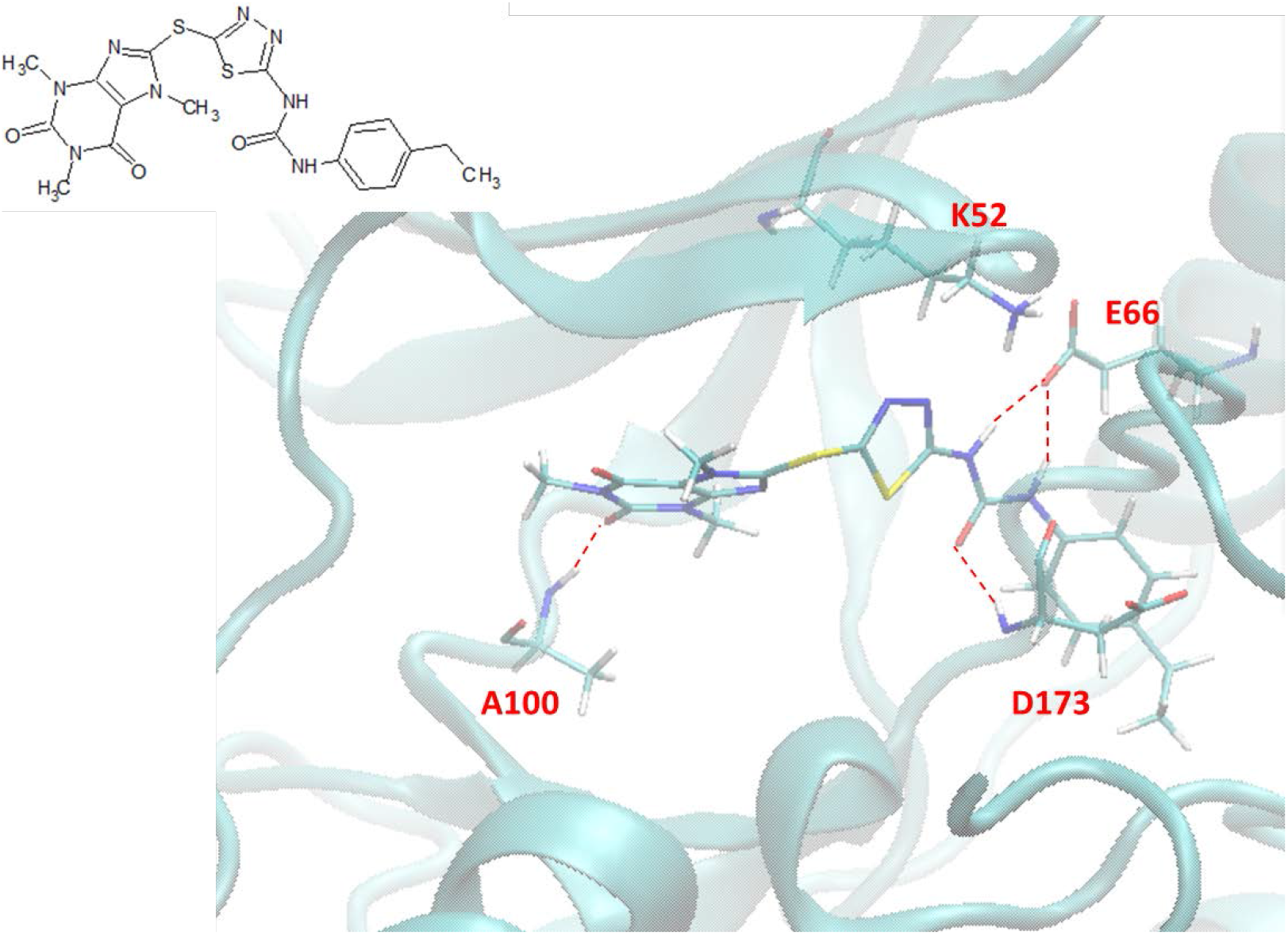
Predicted binding modes of ligand **1** (top) and compound 1 (bottom) by the Superposition and Single-Point Energy Evaluation method. The crystal structure of the reference ligand is colored green for comparison.

We picked 20 compounds from the top ones by the Superposition and Single-Point Energy Evaluation method, plus two selection rules: (1) candidates are able to form at least one hydrogen bond with the hinge region (residues 97 to 100); (2) candidates don’t have toxic substructures. We eye inspected the predicted conformations of the top compounds until the pool was filled up. Table 3 lists the molecular structures of the candidates. When compared with the top 20 compounds ranked by the Superposition and Single-Point Energy Evaluation method, the 20 candidates for VM2 evaluation are more similar to each other. For instance, compounds 1 and 2 are different by only one atom; compounds 4, 21, 37, and 79 share the same scaffold. Similarly, compounds 34 and 119, compounds 36, 46 and 83, compounds 125 and 128 belong to 3 different congeneric series. The VM2 free energy calculation took 3 to 7 days per compound on 4 CPU cores as above.

**Table 3.**
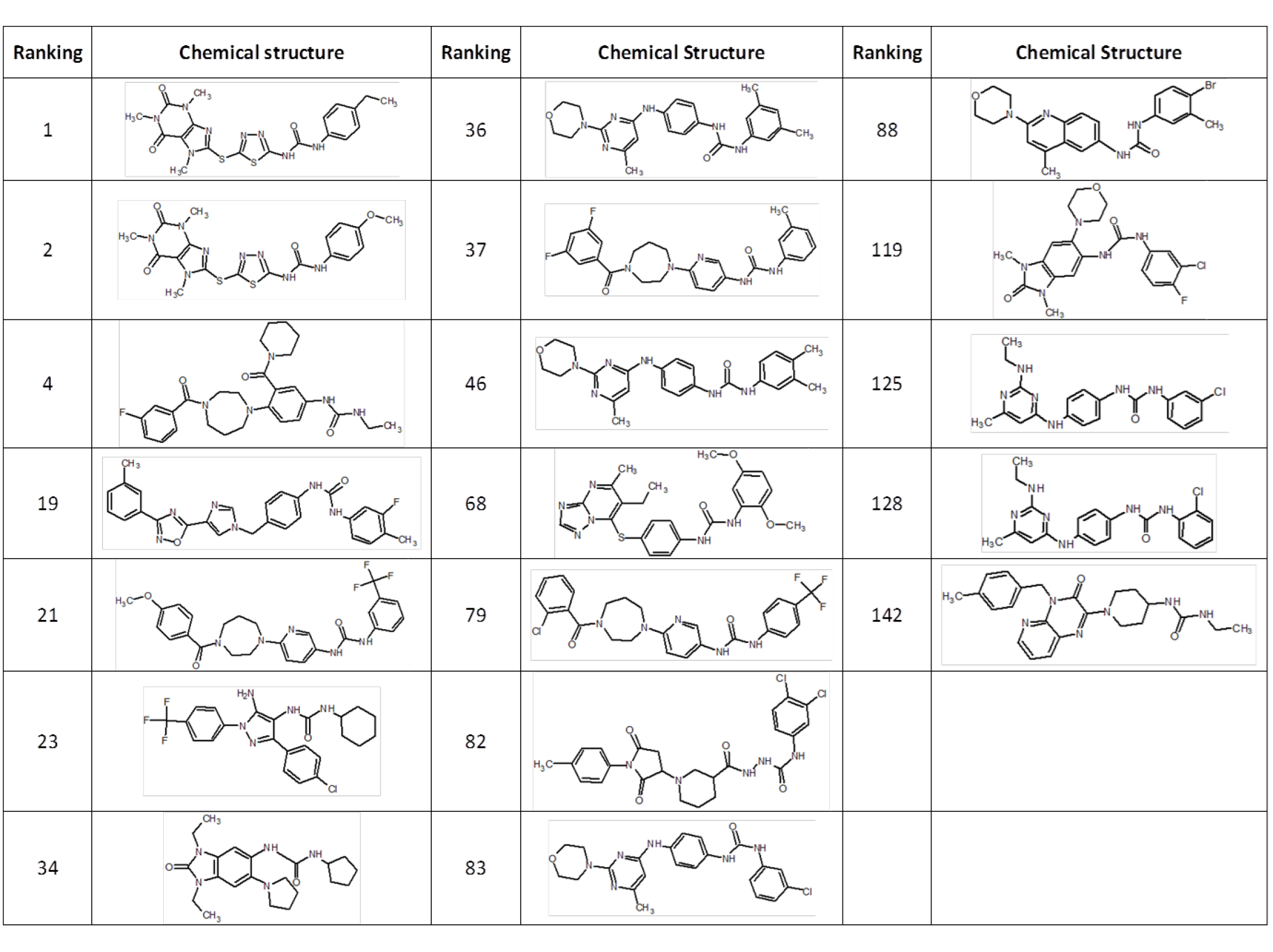
Molecular structures of the 20 candidate compounds picked for VM2 free energy evaluation. The rankings are from the Superposition and Single-Point Energy Evaluation method.

| Ranking | Chemical structure | Ranking | Chemical Structure | Ranking | Chemical Structure |
| --- | --- | --- | --- | --- | --- |
| 1 |  | 36 |  | 88 |  |
| 2 |  | 37 |  | 119 |  |
| 4 |  | 46 |  | 125 |  |
| 19 |  | 68 |  | 128 |  |
| 21 |  | 79 |  | 142 |  |
| 23 |  | 82 |  |  |  |
| 34 |  | 83 |  |  |  |

Table 4 presents the calculated binding free energies (ΔG) and their energy components of the 20 candidates. Only 8 candidates have negative ΔG and none of them have ΔG lower than ligand **1**. Some candidates have a large valence energy term which includes all the bonded interactions. This indicates that these compounds have much internal stress at the protein binding site. This energy term can be considered as an indicator of how well a compound can fit the binding site. The candidates with positive ΔG pay large desolvation penalty and their overall electrostatic energies are consistently greater than 30.0 kcal/mol, about 10.0 kcal/mol higher than those of the candidates with negative ΔG. Compounds 1 and 2 were also ranked as the top 2 compounds by VM2. When compared with ligand **1**, they have much higher (less favorable) ΔE_VDW_ presumably due to their smaller molecular sizes, but they pay less entropy penalty and gain much lower (more favorable) *ΔE_Coulomb_*. Compound 88 was ranked the 3rd by VM2. Its Coulombic energy is on the same level as that of ligand **1** and its van der Waals contribution is similar to that of compounds 1 and 2. It is its reduced desolvation penalty and entropy penalty that boost its calculated binding free energy.

**Table 4.** The calculated free energy and energy components of the 20 candidate compounds by VM2. Please refer to Table 2 for the meaning of each energy term.

| compounds | $\Delta E_{\text{Valence}}$ | $\Delta E_{\text{Coulomb}}$ | $\Delta E_{\text{PB}}$ | $\Delta E_{\text{NP}}$ | $\Delta E_{\text{VDW}}$ | $\Delta H$ | $-T\Delta S$ | $\Delta G_{\text{calc}}$ | $\Delta G_{\text{exp}}$ |
| --- | --- | --- | --- | --- | --- | --- | --- | --- | --- |
| 1 | 2.575 | -37.351 | 59.884 | -6.063 | -66.181 | -47.136 | -28.567 | -18.569 | -10.116 |
| 2 | 5.835 | -30.900 | 51.406 | -5.825 | -69.650 | -49.134 | -31.329 | -17.805 | -9.528 |
| 88 | 5.437 | -20.290 | 46.567 | -5.697 | -68.374 | -42.357 | -25.080 | -17.277 | -6.784 |
| 21 | 12.007 | -93.269 | 109.888 | -7.106 | -71.414 | -49.894 | -34.696 | -15.198 |  |
| 37 | 8.164 | -88.559 | 110.296 | -6.768 | -62.763 | -39.630 | -29.486 | -10.144 |  |
| 79 | 15.828 | -90.286 | 106.577 | -6.888 | -62.442 | -37.211 | -28.617 | -8.594 |  |
| 68 | 8.595 | -19.085 | 47.124 | -6.280 | -65.626 | -35.272 | -28.657 | -6.615 |  |
| 125 | 5.678 | -17.151 | 38.969 | -5.712 | -55.414 | -33.630 | -32.058 | -1.572 |  |
| 128 | 0.682 | -11.092 | 43.001 | -5.845 | -55.046 | -28.300 | -31.304 | 3.004 |  |
| 83 | 10.269 | -9.602 | 42.038 | -6.105 | -63.284 | -26.684 | -31.977 | 5.293 |  |
| 36 | 11.726 | -9.625 | 43.528 | -6.369 | -65.677 | -26.417 | -32.157 | 5.740 |  |
| 46 | 11.314 | -14.672 | 50.305 | -6.328 | -64.128 | -23.509 | -31.125 | 7.616 |  |
| 119 | 5.942 | -11.856 | 49.080 | -5.776 | -58.916 | -21.526 | -30.401 | 8.875 |  |
| 142 | 6.350 | -8.376 | 42.859 | -5.845 | -53.171 | -18.183 | -29.681 | 11.498 |  |
| 34 | 16.957 | -10.557 | 40.449 | -5.221 | -63.213 | -21.585 | -35.780 | 14.195 |  |
| 4 | 13.136 | -10.238 | 45.821 | -6.210 | -58.802 | -16.293 | -30.728 | 14.435 |  |
| 23 | 7.329 | -22.129 | 66.030 | -6.459 | -58.193 | -13.422 | -32.146 | 18.724 |  |
| 19 | 14.920 | -16.982 | 50.464 | -6.551 | -57.835 | -15.984 | -37.405 | 21.421 |  |
| 82 | 62.593 | -5.770 | 58.124 | -6.423 | -29.028 | 79.496 | -42.367 | 121.863 |  |
| reference | 1.031 | -19.845 | 60.764 | -8.548 | -93.837 | -60.435 | -39.487 | -20.948 | -10.324 |

The lowest energy conformation of compound 1 with CDK8 (Figure 6) predicted by VM2 is similar to the one predicted by the Superposition and Single-Point Energy Evaluation method. The key hydrogen bonds with E66, D173 and A100 are well kept. Furthermore, its 1,3,4-thiadiazole forms H bonding with K52. This may explain its favorable Coulombic contribution. Its back bone is full of aromatic rings, which make the whole molecular structure relatively rigid and reduce the entropy penalty. The extended conformation helps the bulky aromatic rings on both ends stay at the hydrophobic deep pocket and front pocket respectively. They are surrounded by nonpolar residues and make extensive van der Waals interactions with them. Compound 1 has a relatively small molecular weight (MW), 473 daltons. As a comparison, the MW of ligand **1** is 640 daltons. Compound 2 is different from compound 1 by only one atom, and it has the exactly same binding mode as compound 1. The predicted lowest energy conformation of compound 88 can be found in Figure S4. The starting conformation of compound 88 for the VM2 free energy calculation has H bonding with A100. But VM2 ended up with the morpholine ring rotating by 90° and losing the contact with A100. Compound 88 has a bulky quinoline ring in the middle of its backbone. This helps to reduce its entropy penalty, but also limits the arrangement of the whole molecular structure to accommodate H bonding between the morpholine ring and A100.

**Figure 6.**
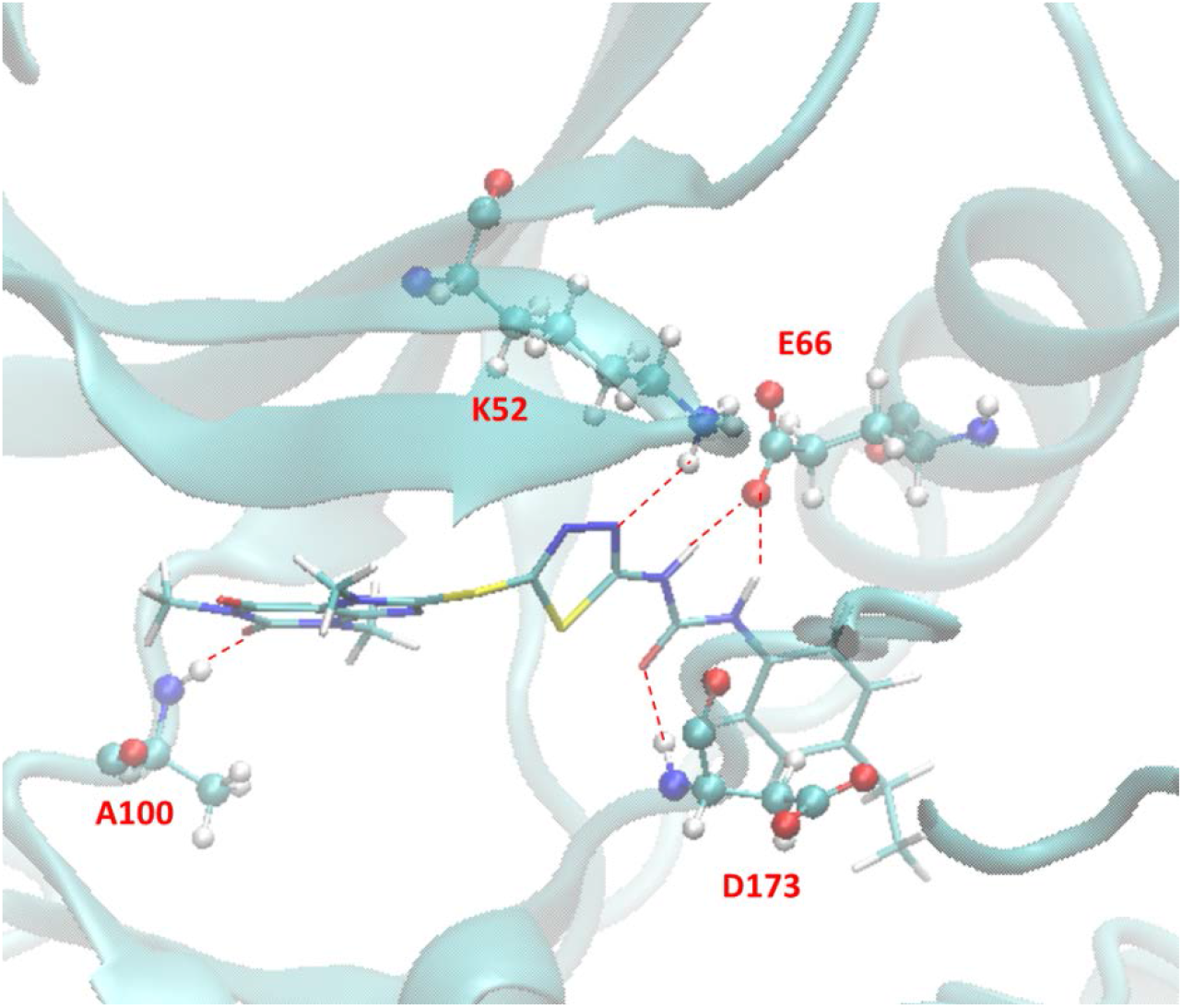
Predicted lowest energy conformation of compound 1 with CDK8 by VM2.

We purchased the top 3 candidates ranked by VM2, i.e. compounds 1, 2 and 88, and had bio-assay testing with them. The result (*ΔG_exp_*) can be found in Table 4. Compounds 1 and 2 show strong binding affinity with CDK8, the K_d_ values of which are 42.5 nM and 114 nM respectively. The K_d_ value of compound 88 is 11.4 μM, which is barely lower than the no-binding cutoff (20 μM) in the testing. Both the Superposition and Single-Point Energy Evaluation method and the VM2 method ranked the 3 compounds correctly. Compound 1 achieved a binding affinity comparable to those of the two most potent reference ligands (K_d_ values of 10 nM and 30 nM for ligand **2** and ligand **1** respectively). It has a drug-like structure with dimethyl-xanthine, 1,3,4-thiadiazole and benzene moieties on its backbone. Figure 7 presents the correlation between the calculated and experimental free energies when the data points of the 3 candidates are plotted together with the reference ligands. The correlation coefficient drops from 0.71 to 0.47 after these data points are added. The poorer correlation is mainly caused by the overestimated binding free energy of compound 88. If this data point is removed from the plot, the correlation coefficient becomes 0.72. Despite the overestimation, VM2 predicts that compound 88 have no contact with the hinge region, which is probably correct and explains the weak affinity of this compound. The interaction with the hinge region is important to ligand binding to CDK8 and is found in the binding modes of the two strong binders, compounds 1 and 2.

**Figure 7.**
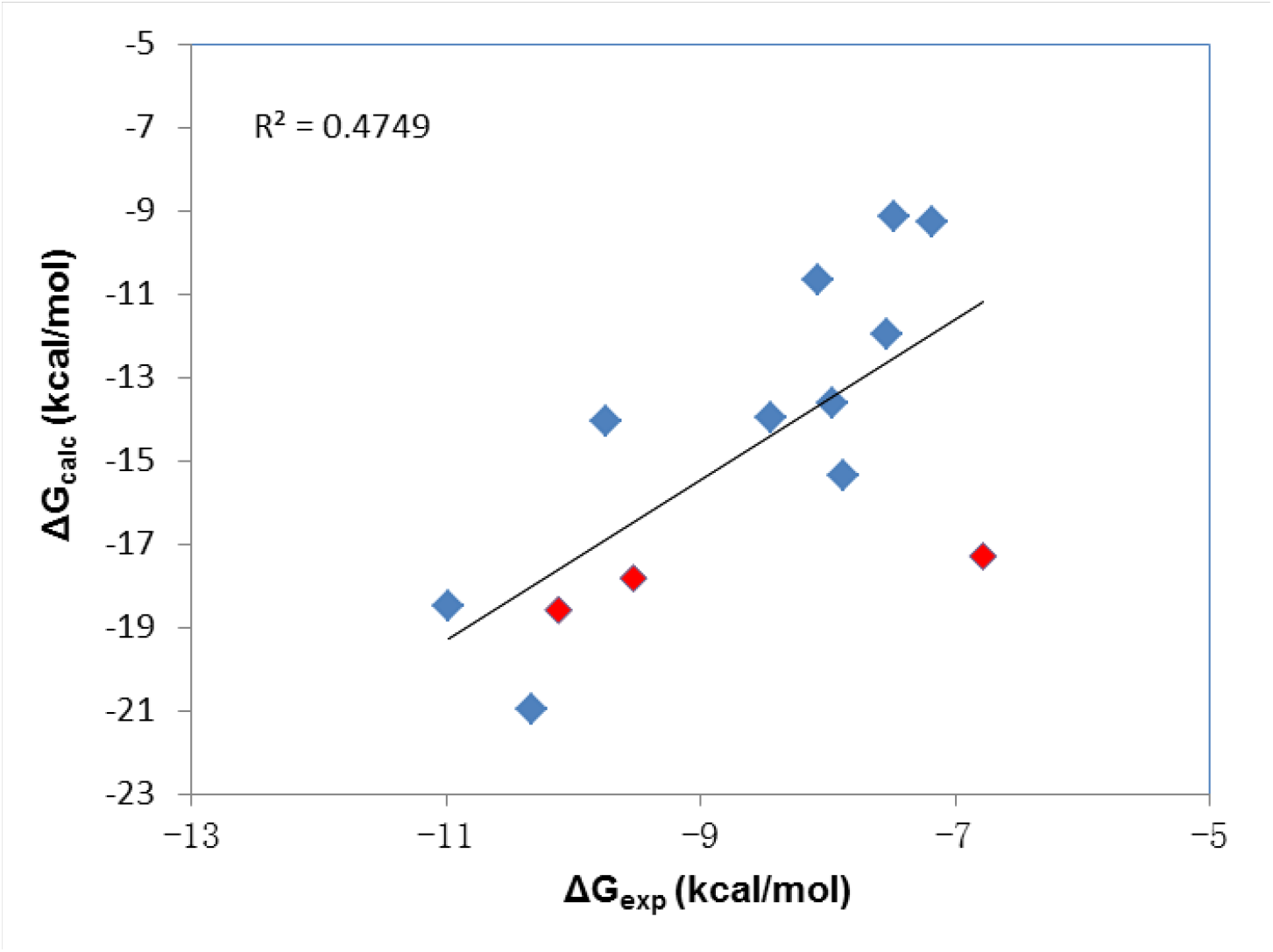
The correlation between the experimental and calculated binding free Energies for the 11 reference ligands and 3 tested compounds. Data points for the reference ligands and tested compounds are colored blue and red respectively.

With our virtual drug screening package we successfully discovered a new potent CDK8 type II ligand, compound 1, which is comparable to the published reference ligands. It is worth to mention that the whole virtual screening process was finished in ten days. The effectiveness of this novel method is originated from the combination of two powerful computational methods and the knowledge learned from the thermodynamics analysis on reference ligands. The Superposition and Single-Point Energy Evaluation method was able to efficiently estimate the binding poses for candidate compounds with a co-crystal structure as reference. The binding energies obtained by this method are based upon force field energy equations and PB solvation model. Energy minimization is implemented on candidate compounds and binding site residues until convergence. Therefore, this method can offer high accuracy in energy prediction while maintaining relatively high speed. The VM2 free energy calculation method carries out thorough conformation sampling and considers both enthalpy and entropy contributions to free energies. This method is fast and accurate enough to be used as the last check point in virtual drug screening. And since it provides not only the total free energy and its components but also the energy contributions from the portions of the molecular systems, it is a useful tool for ligand optimization as well.

This novel virtual drug screening tool is also valuable in the context of fragment-based drug discovery. The structural moieties that make key interactions with a target protein in existing co-crystal structures can be employed separately in similarity or substructure database searches for fragments, which can then be merged or linked together to generate new ligands. Fragments generally have weak affinities with target proteins and pose significant challenges for screening through biophysical techniques [50, 51]. The two energy evaluation methods in the virtual drug screening tool have the accuracy to solve the problem of fragment screening.

## 4. Conclusion

We developed a novel virtual drug screening package and applied it to the discovery of new CDK8 type II ligands. The core of this method consists of two energy evaluation methods: Superposition and Single-Point Energy Evaluation, and VM2 free energy calculation. They are able to efficiently and accurately estimate the binding energies of candidate compounds. In this research work we analyzed binding free energies and the energy components of 11 reference CDK8 type II ligands with VM2, and extracted the information which was proved helpful for virtual drug screening.

The VM2 method accurately predicted the binding modes for the reference ligands and their RMSDs are all less than 2.0 Å for the ligand atoms and atoms of binding site residues. The correlation coefficient is 0.71 between the calculated and measured free energies. The free energy and MD calculations successfully revealed the factors that play important roles in the ligand binding with CDK8 DMG-out conformation. The overall driven force of the binding is van der Waals interaction, but for some ligands Coulombic energy is also important to make the binding to occur. The urea moiety contributes the majority of H bonding between the reference ligands and CDK8 and acts as the anchorage to stabilize the ligands. The analysis on the strong binders also suggests that there is room for all of them to improve their binding affinities.

Starting with the urea moiety, we implemented the virtual drug screening package and singled out three compounds for bio-assay testing. The ranking from the experimental result is completely consistent with the predicted rankings by both Superposition and Single-Point Energy Evaluation method and VM2 free energy calculation method for the three compounds. We successfully discovered a new potent drug-like compound with a K_d_ value of 42.5 nM. This compound was ranked #1 by both methods. Interestingly, top 2 compounds are different by only one atom but have a nearly 3-fold difference in binding affinity. This was accurately predicted by both energy evaluation methods. Therefore, our novel virtual drug screening package is accurate and efficient enough to be used in drug design projects. We believe this work has significant impact to the field of drug discovery.

## Acknowledgement

This study was supported by the US National Institutes of Health (GM-109045).

## Supplemental Materials

**Figure S1.**
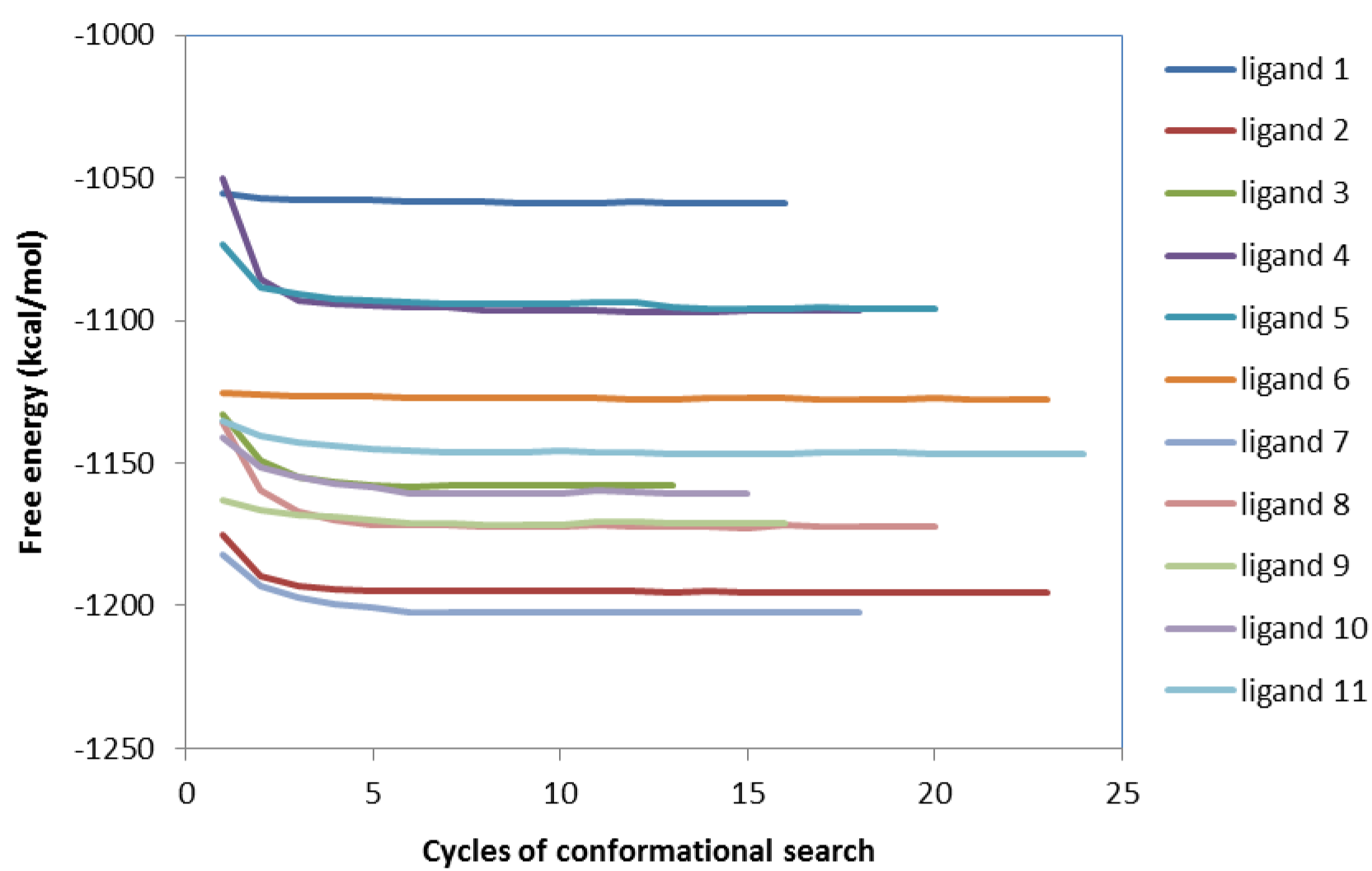
Convergence of computed free energies of protein-ligand complexes of the 11 reference ligands, as a function of the number of search cycles.

**Figure S2.**
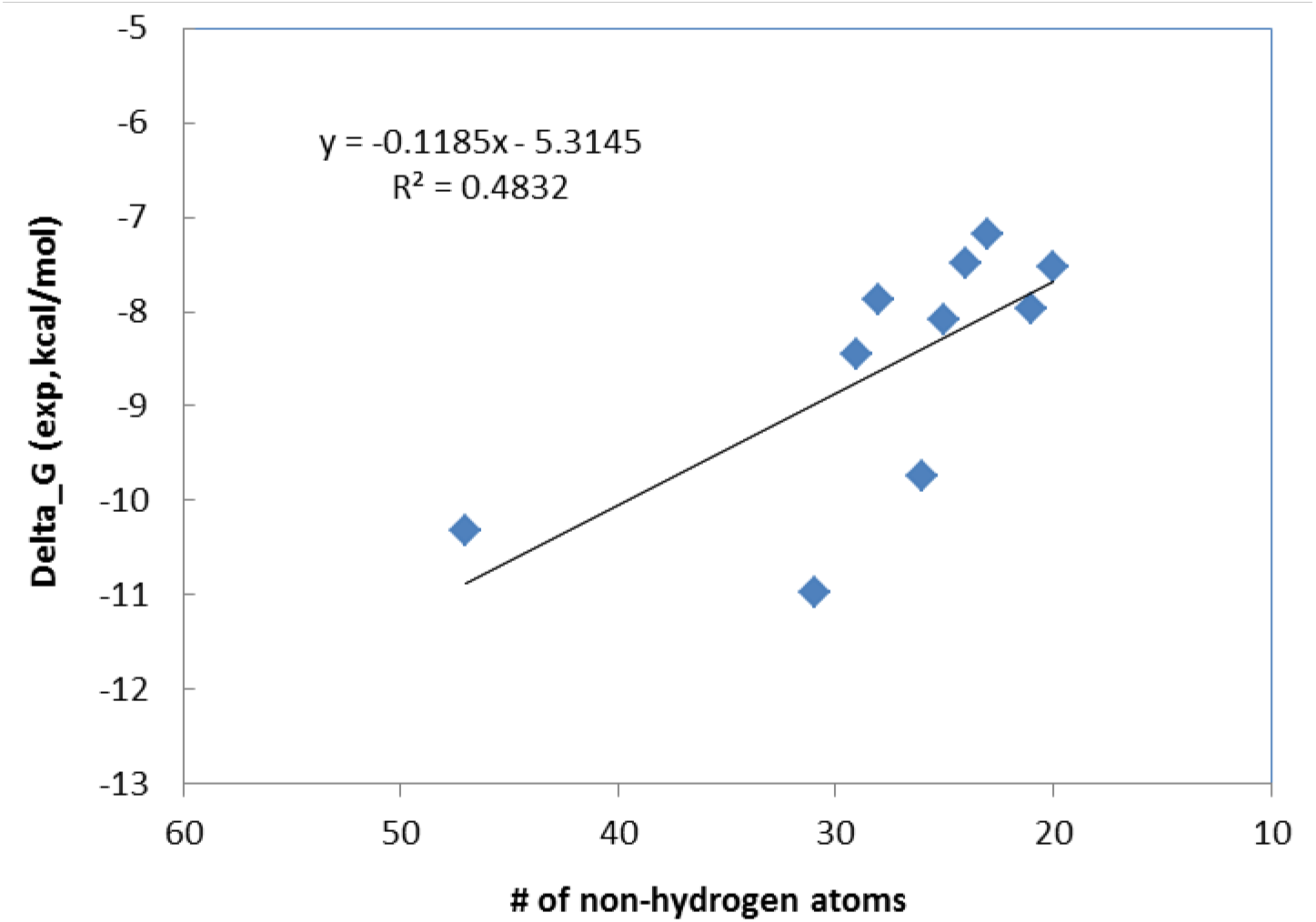
The correlation of the number of non-hydrogen atoms of ligands with the experimental binding free energies for the 11 reference ligands.

**Figure S3.**
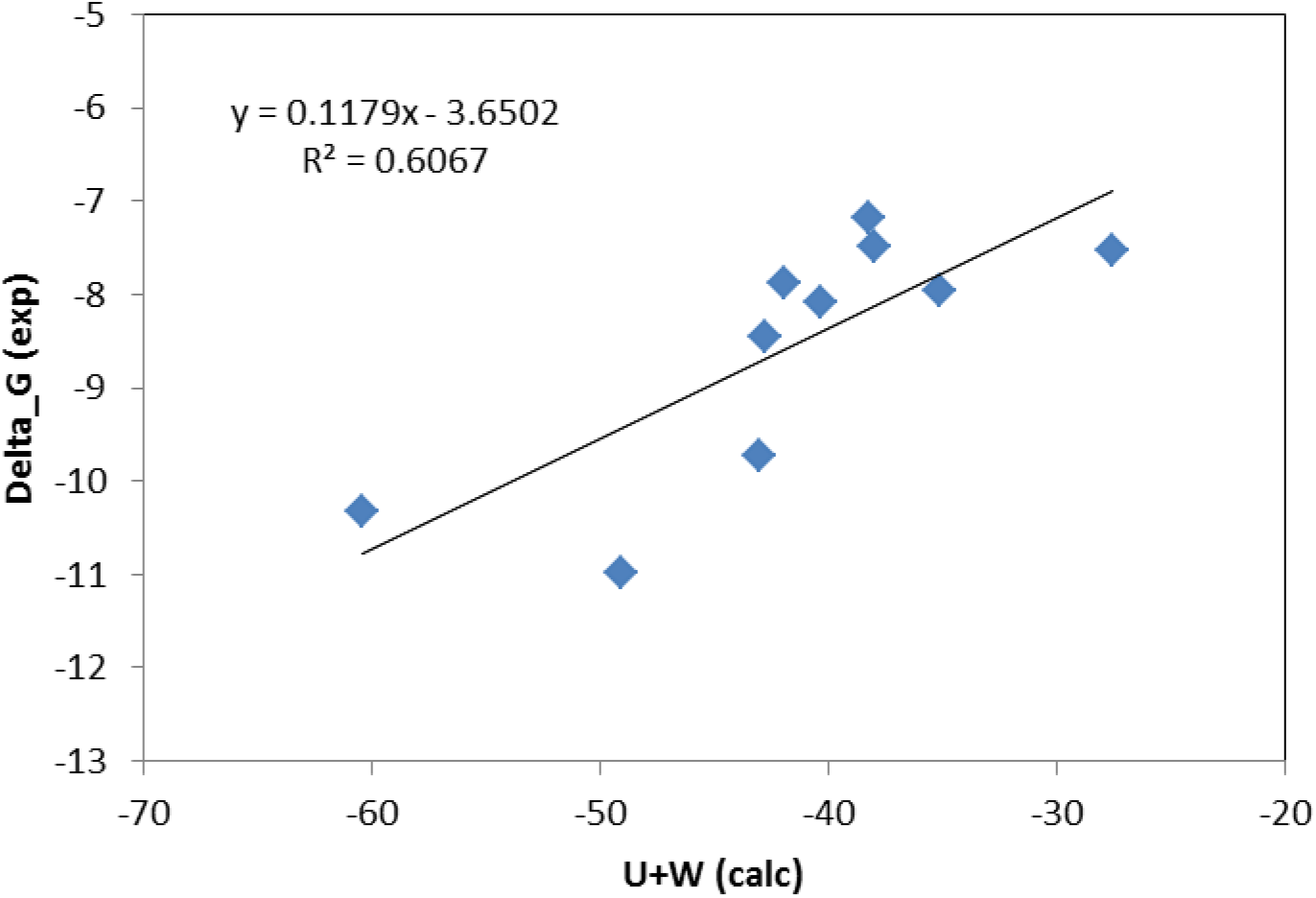
The correlation of the calculated enthalpy with the experimental binding free energies for the 11 reference ligands.

**Figure S4.**
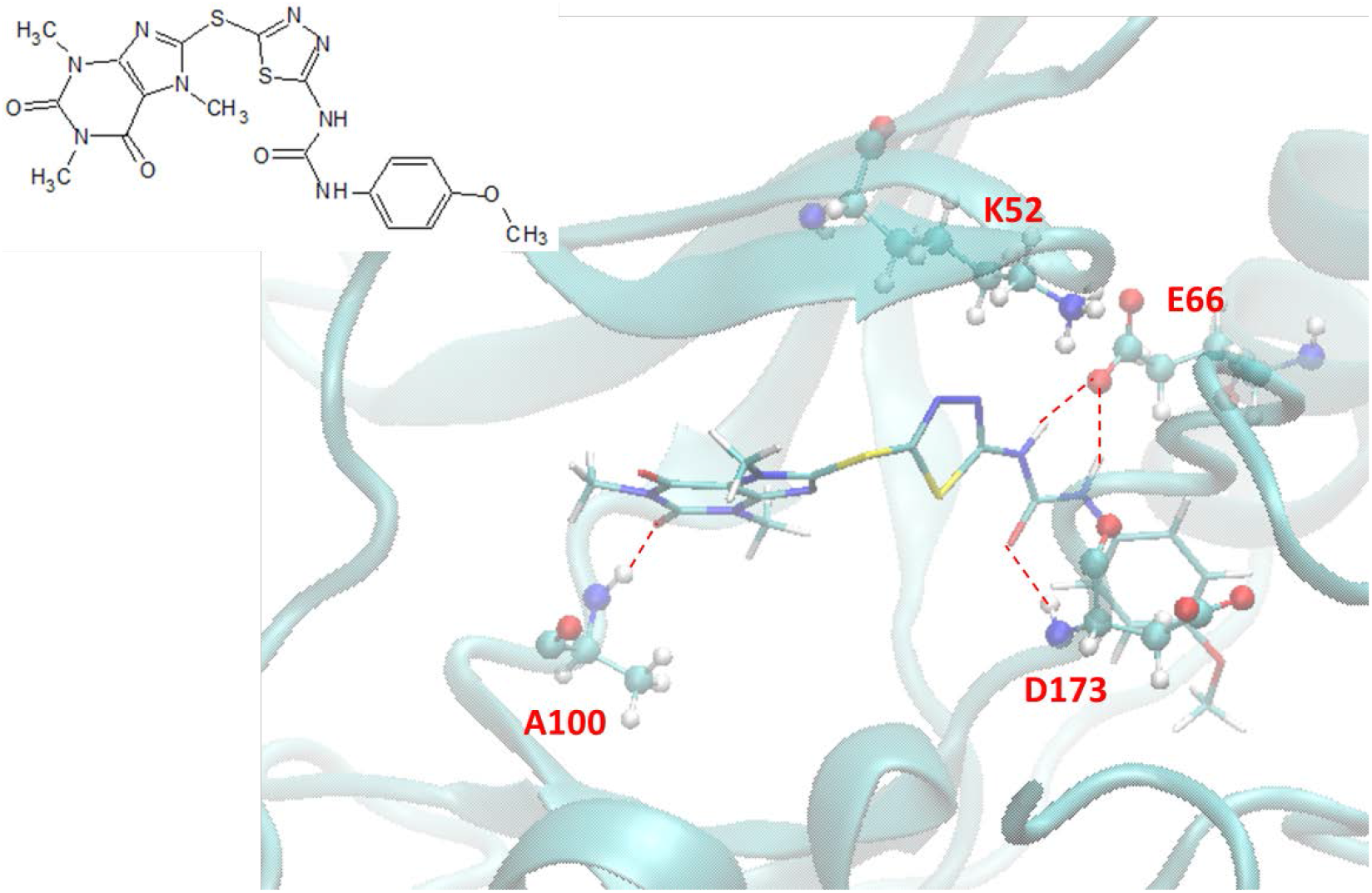
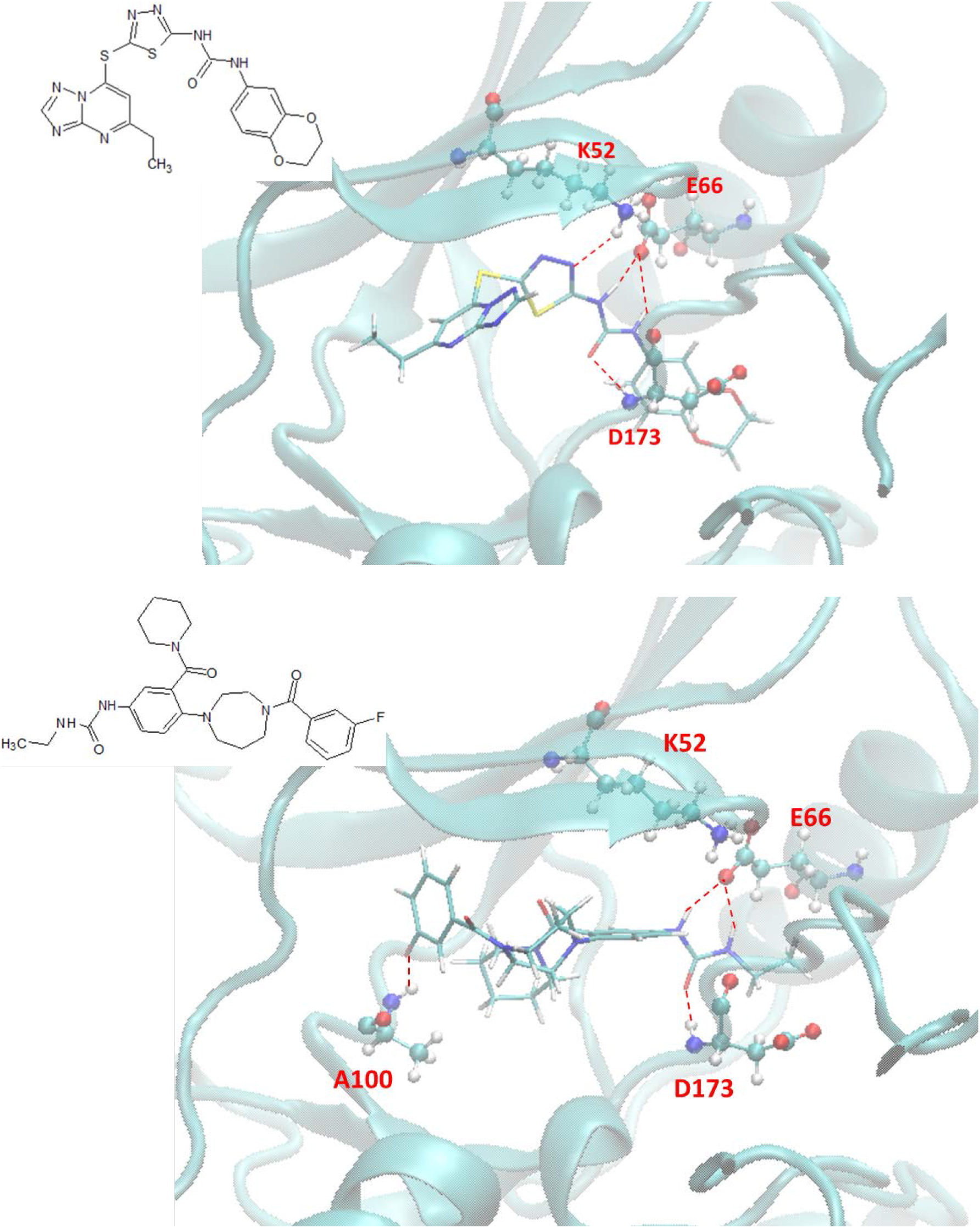
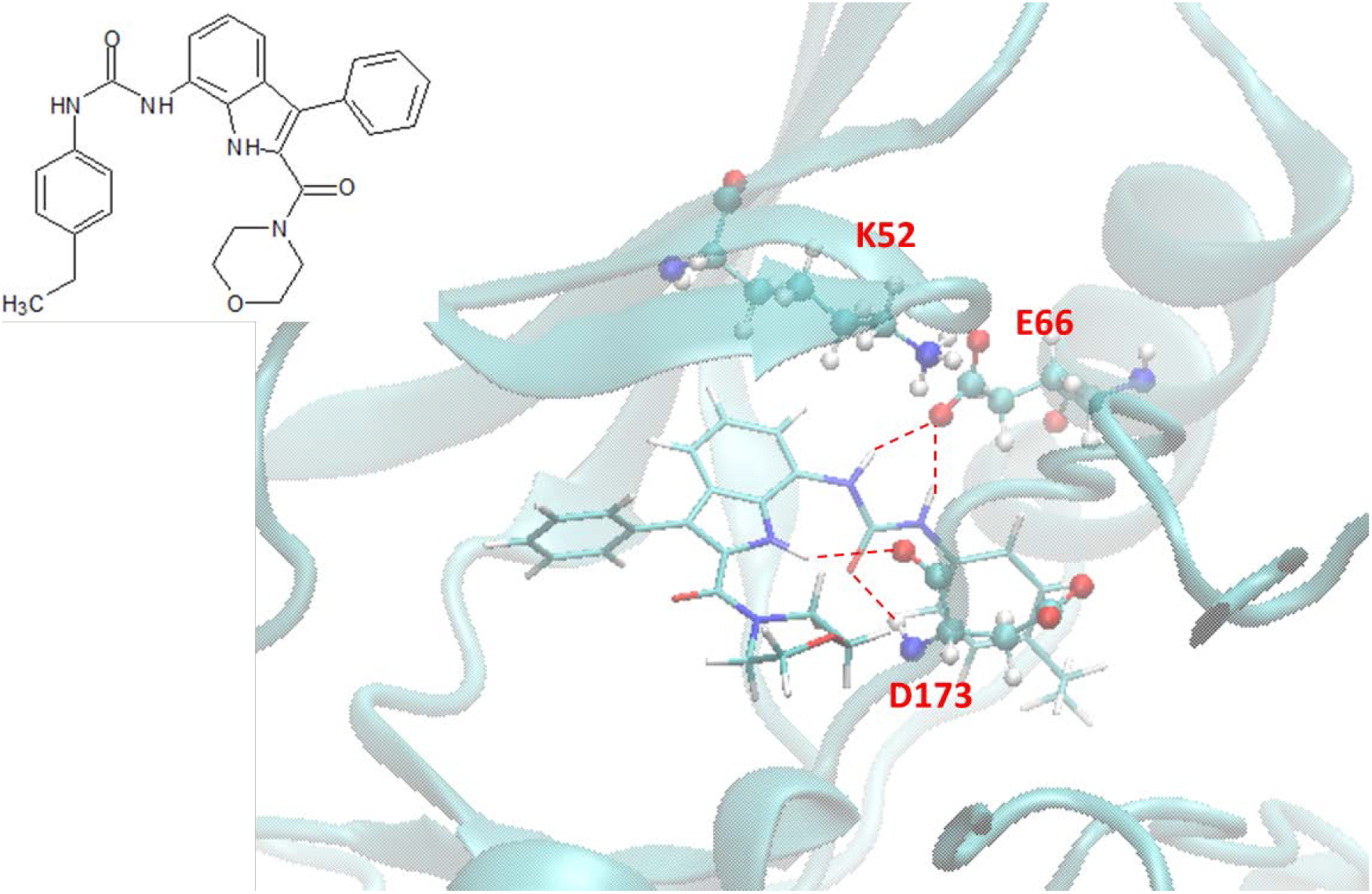
Predicted binding modes of compounds 2, 3, 4 and 5 with CDK8 (from top to bottom) by the Superposition and Single-Point Energy Evaluation method.

**Figure S5.**
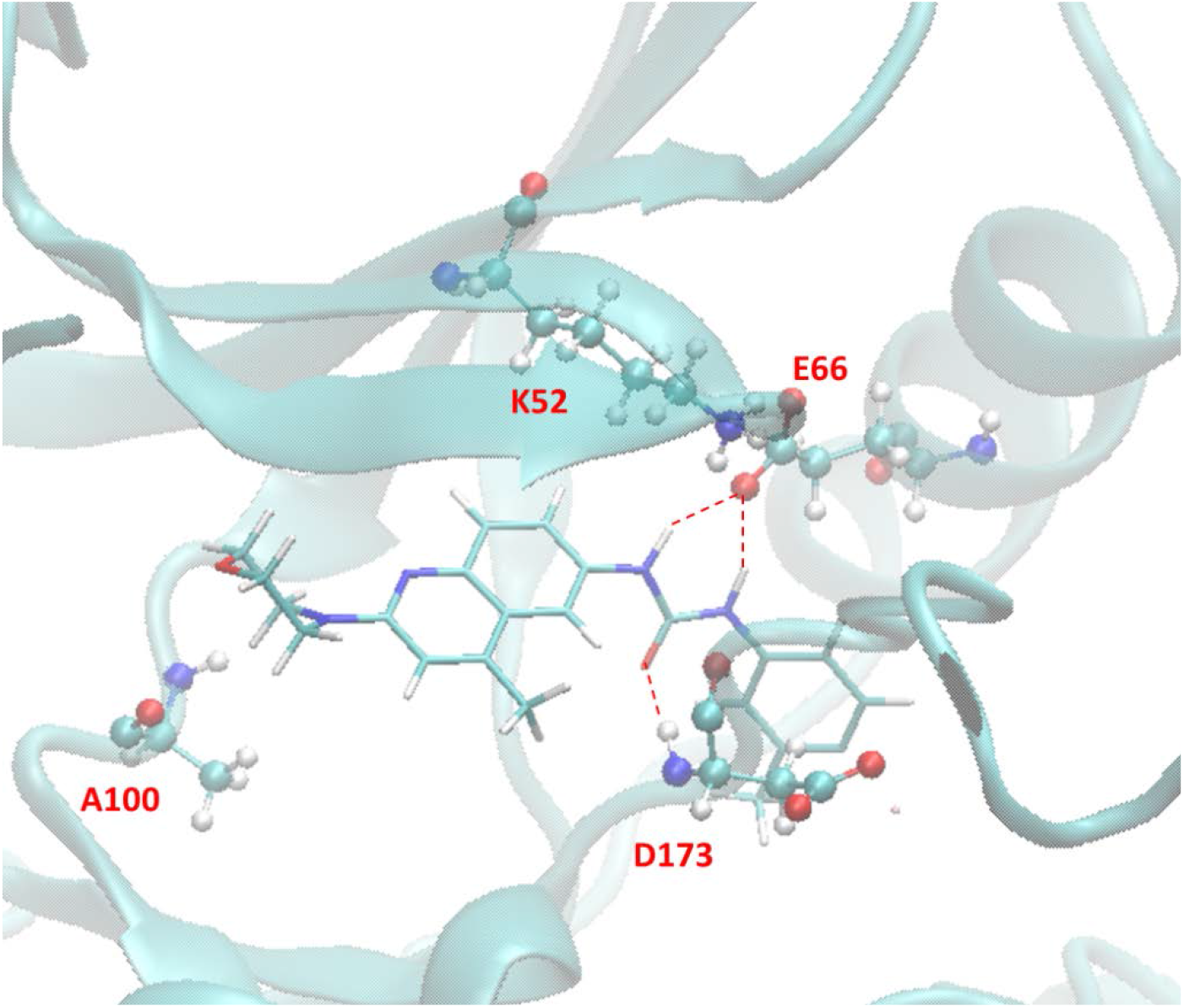
The predicted lowest energy conformation of compound 88 with CDK8 by VM2.

**Table S1.** Top 100 compounds ranked by the Superposition and Single-Point Energy Evaluation method. The unit is kcal/mol. The reference ligand (ligand **1**) has a binding energy of –75.104 kcal/mol.

|  |  |  |  |  |  |  |  |
| --- | --- | --- | --- | --- | --- | --- | --- |
| 1 | -63.937 | 26 | -55.175 | 51 | -53.562 | 76 | -52.827 |
| 2 | -61.274 | 27 | -55.141 | 52 | -53.538 | 77 | -52.816 |
| 3 | -58.569 | 28 | -55.133 | 53 | -53.523 | 78 | -52.683 |
| 4 | -58.214 | 29 | -55.06 | 54 | -53.511 | 79 | -52.65 |
| 5 | -57.632 | 30 | -54.967 | 55 | -53.432 | 80 | -52.644 |
| 6 | -57.476 | 31 | -54.909 | 56 | -53.313 | 81 | -52.586 |
| 7 | -57.455 | 32 | -54.769 | 57 | -53.29 | 82 | -52.569 |
| 8 | -57.29 | 33 | -54.608 | 58 | -53.22 | 83 | -52.52 |
| 9 | -57.144 | 34 | -54.588 | 59 | -53.198 | 84 | -52.506 |
| 10 | -57.135 | 35 | -54.542 | 60 | -53.19 | 85 | -52.487 |
| 11 | -56.791 | 36 | -54.49 | 61 | -53.183 | 86 | -52.486 |
| 12 | -56.756 | 37 | -54.458 | 62 | -53.178 | 87 | -52.361 |
| 13 | -56.49 | 38 | -54.433 | 63 | -53.125 | 88 | -52.358 |
| 14 | -56.398 | 39 | -54.364 | 64 | -53.118 | 89 | -52.352 |
| 15 | -55.929 | 40 | -54.182 | 65 | -53.114 | 90 | -52.301 |
| 16 | -55.916 | 41 | -54.033 | 66 | -53.103 | 91 | -52.271 |
| 17 | -55.909 | 42 | -54.013 | 67 | -53.101 | 92 | -52.269 |
| 18 | -55.847 | 43 | -53.996 | 68 | -53.076 | 93 | -52.2 |
| 19 | -55.767 | 44 | -53.962 | 69 | -53.064 | 94 | -52.168 |
| 20 | -55.708 | 45 | -53.868 | 70 | -53.035 | 95 | -52.164 |
| 21 | -55.59 | 46 | -53.864 | 71 | -53.027 | 96 | -52.143 |
| 22 | -55.525 | 47 | -53.843 | 72 | -52.927 | 97 | -52.119 |
| 23 | -55.441 | 48 | -53.716 | 73 | -52.894 | 98 | -52.107 |
| 24 | -55.272 | 49 | -53.68 | 74 | -52.848 | 99 | -52.09 |
| 25 | -55.269 | 50 | -53.639 | 75 | -52.836 | 100 | -52.077 |

**Table S2.**
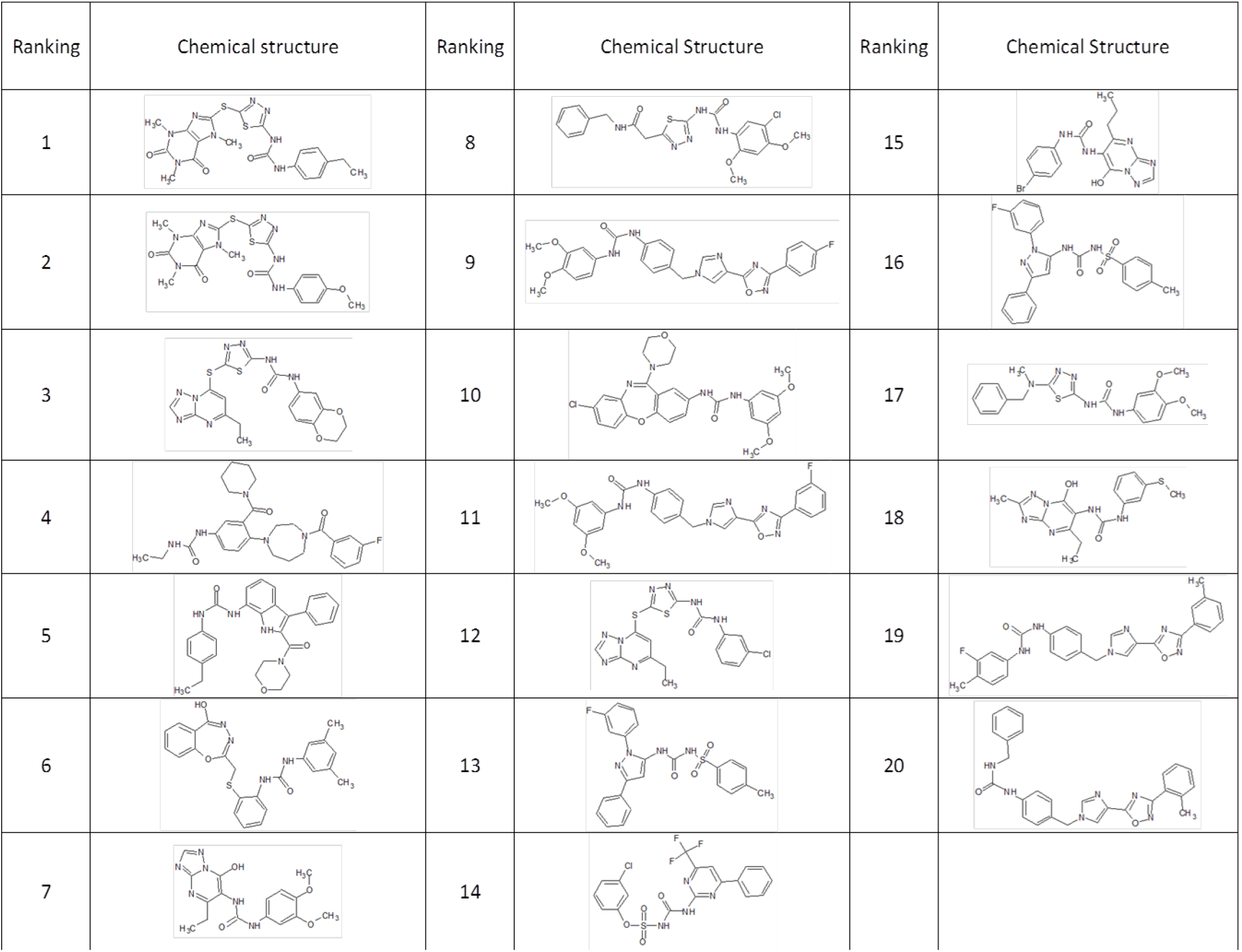
Molecular structures of top 20 compounds ranked by the Superposition and Single-Point Energy Evaluation method.

| Ranking | Chemical structure | Ranking | Chemical Structure | Ranking | Chemical Structure |
| --- | --- | --- | --- | --- | --- |
| 1 |  | 8 |  | 15 |  |
| 2 |  | 9 |  | 16 |  |
| 3 |  | 10 |  | 17 |  |
| 4 |  | 11 |  | 18 |  |
| 5 |  | 12 |  | 19 |  |
| 6 |  | 13 |  | 20 |  |
| 7 |  | 14 |  |  |  |

